# The Alternate Ligand Jagged Enhances the Robustness of Notch Signaling Patterns

**DOI:** 10.1101/2022.11.16.516674

**Authors:** Mrinmoy Mukherjee, Herbert Levine

**Affiliations:** Center for Theoretical Biological Physics, Northeastern University, Boston, MA; Depts. of Physics and Bioengineering, Northeastern University, Boston, MA

## Abstract

The Notch pathway, an example of juxtacrine signaling, is an evolutionary conserved cell-cell communication mechanism. It governs emergent spatiotemporal patterning in tissues during development, wound healing and tumorigenesis. Communication occurs when Notch receptors of one cell bind to either of its ligands, Delta/Jagged of neighboring cell. In general, Delta-mediated signaling drives neighboring cells to have an opposite fate (lateral inhibition) whereas Jagged-mediated signaling drives cells to maintain similar fates (lateral induction). Here, By deriving and solving a reduced set of 12 coupled ordinary differential equations for Notch-Delta-Jagged system on a hexagonal grid of cells, we determine the allowed states across different parameter sets. We also show that Jagged (at low dose) acts synergistically with Delta to enable more robust pattern formation, despite of its lateral induction property; this effect is due to competition with Delta over binding with Notch, as experimentally observed in the case of chick inner ear development. Finally, we show that how Jagged can help to expand the bistable (both Uniform and Hexagon phases are stable) region, where a local perturbation can spread over time in a ordered manner to create a biologically relevant, defect-free lateral inhibition pattern.

## I. INTRODUCTION

Notch signaling plays a crucial role in controlling cell-fate decisions during embryonic development.[1, 2] The signaling cascade is initiated via ligand binding to Notch transmembrane receptors, leading to the release of the Notch Intercellular Domain (NICD) and downstream regulation by NICD of its target genes.[1, 3–5] This simple mechanism regulates cell-fate differentiation in different biological system ranging from development of the inner ear,[6, 7] vascular smooth muscle cell development,[5] Drosophila wing disk formation,[1] bristle patterning[5] and cancer metastasis.[8–11]

There are two types of ligands, Delta-like and Jagged-like, which can bind to the Notch receptors on the surface of a neighboring cell, as examples of juxtacrine signaling. The signal can introduce a biochemical feedback between neighboring cells coordinating their cell-fate, which leads to the spatiotemporal pattering in multicellular systems. Notch-ligand binding can also happen in same cell (cis-coupling) apart from the usual interaction between neighboring cells (trans-coupling).[12, 13]

In general, Delta-mediated Notch signaling drives neighboring cells to have an opposite fate, which create an alternate ‘salt and pepper’ pattern of Sender (high ligand, low receptor) and Receiver (low ligand, high receptor) cells in tissue; this is referred to as lateral inhibition. Alternatively, Jagged promotes the similar cell-fate in neighboring cells, giving rise to lateral induction. The full Notch-Delta-Jagged system can act as a three-way switch, giving rise to an additional hybrid state (medium ligand, medium receptor).[14, 15] Also, at an intermediate production rate, adding Jagged to the pure Notch-Delta system can alter the accuracy[16] and robustness[7] of the patterns in various developmental processes.

In this paper, we study the pattern formation problem in the Notch-Delta-Jagged system, extending the framework[17] previously used for the Notch-Delta system. Inspired by the general tissue structure in epithelial monolayers which roughly form a hexagonal lattice,[8] we evaluate the dynamics of Notch, Delta, Jagged and NICD on a 2d hexagonal array of cells. We calculate the phase space across different parameters for which stable (ordered) solutions exist. We mainly focus on the dose-dependent role of Jagged and how it can affect the accuracy and robustness of disordered patterns, generated from uniform (with small noise) initial conditions. In the end, we discuss how an expanded region of bistability (where both Uniform and Hexagon stable phases coexist) can arise in the presence of Notch-Jagged signaling and help to form perfectly ordered patterns. This can be a useful strategy to obtain accurate systems in noisy biological systems.

## II. MODEL

Here, we study the Notch-Delta-Jagged system on a 2d hexagonal array of cells. To incorporate both the basic features of Notch-Delta signaling induced lateral inhibition pattern (Fig. 1a) and Notch-Jagged signaling induced lateral induction pattern (Fig. 1b), we use the following deterministic ordinary differential equations (ODEs)[14, 15] based on previously introduced models,[13, 18–20] involving the concentrations of Notch (*N*), Delta (*D*), Jagged (*J*) and NICD (*I*),

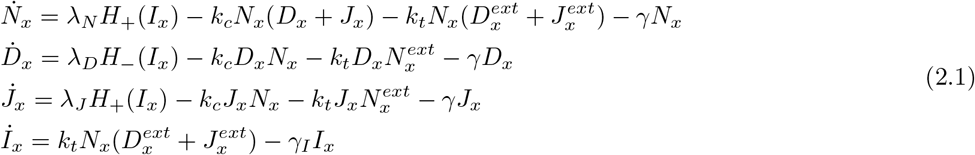

where, *x* refer to the positions of the cells on hexagonal lattice. 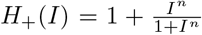 and 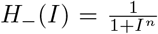 are the Hill functions to represent the effect of NICD (*I*) (via transcriptional regulation) on the production rate of Notch (*N*), Delta (*D*) and Jagged (*J*). *k_c_* and *k_t_* are the strengths of cis-inhibition and trans-activation respectively. λ_*N*_, λ_*D*_ and λ_*J*_ are the production rates of *N*, *D* and *J* respectively. *γ* represents the degradation rate of *N*, *D*, *J* (assumed equal) and *γ_I_* the degradation rate of *I*. (*N*, *D*, *J*)^*ext*^ refers to the average over the 6 nearest neighbors of cell *x*.

**FIG. 1:**
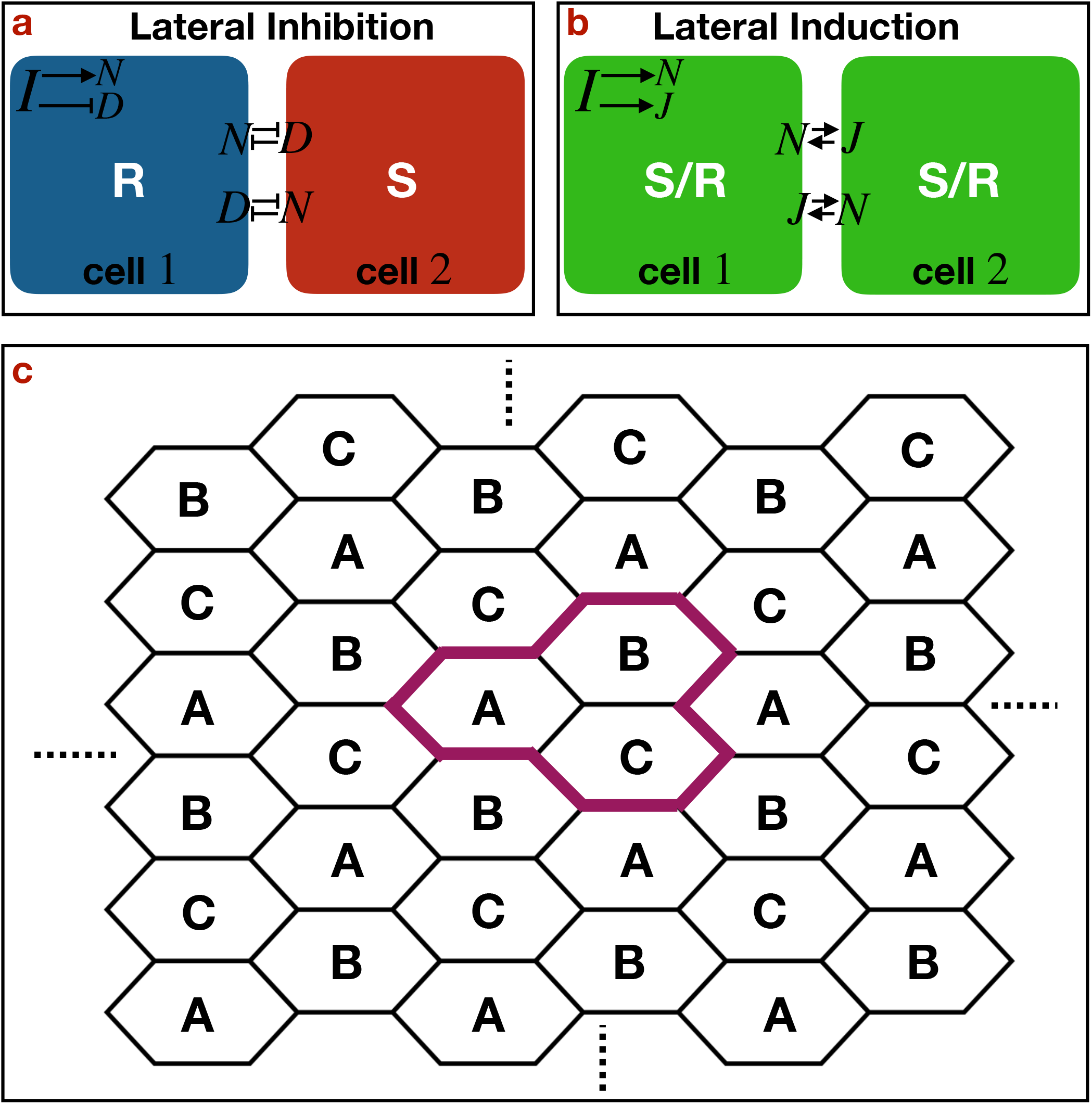
Schematics of lateral inhibition and lateral induction. Schematic diagram of (a) lateral inhibition between two neighboring cells, (b) lateral induction between two neighboring cells and (c) a hexagonal lattice system spanned by A-B-C unit cells.

We use a typical set of parameters taken from the literature[13–15, 17]: *k_c_* = 0.1, *k_t_* = 0.04, *γ* = 0.1, *γ_I_* = 0.5, the Hill coefficients for Notch and Delta (*n_N_* = *n_D_* = 2) and for Jagged (*n_J_* = 5) and vary λ_*N*_, λ_*D*_ and λ_*J*_. We also investigate the effect of changing *k_c_* and *k_t_*. All the parameters used in the Figures throughout the manuscript are listed in Table S1 (see SI).

We are interested in hexagonal ordered patterns. These patterns on a hexagonal lattice are invariant under the translation with vectors 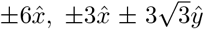 (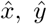 are the unit vectors along the axes (*xy*) and unit length is 1/2 the length of the hexagonal sides). Thus, the concentrations of Notch (*N*), Delta (*D*), Jagged (*J*) and NICD(*I*) everywhere on the lattice are completely determined by their concentrations on the cells labelled by A, B and C (Fig. 1c). Thereby, the entire problem (Eq. 2.1) of hexagonal ordered patterns for a multicellular system is reduced to 12 coupled ODEs (explicit eqns. given in the SI).

## III. RESULTS AND DISCUSSION

### A. Phase diagrams

We solve the reduced set of ODEs numerically to find the parts of parameter space for which the uniform state ((*N*,*D*,*J*,*I*)_*A*_ = (*N*,*D*,*J*,*I*)_*B*_ = (*N*,*D*,*J*,*I*)_*C*_) is unstable with respect to perturbations. First, we find the fixed points where 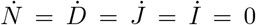 and analyze their stability via linear stability analysis across the parameter space. For a fixed value of λ_*N*_ and λ_*J*_ the uniform solution becomes unstable for 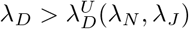 via a transcritical bifurcation (Fig. 2) and overlaps non-uniform solutions where the concentrations on two sublatices (say B, C) are always identical, differing from the concentrations on the remaining sublatice A, such that, (*N*,*D*,*J*,*I*)_*B*_ = (*N*,*D*,*J*,*I*)_*C*_ ≠ (*N*,*D*,*J*,*I*)_*A*_. There are two types of these hexagonal solution; ‘hexagon’ (high D cells surrounded by high N cells), and ‘antihexagon’ (high N cells surrounded by high D cells). Labeling the high *D* cells as ‘Senders’ (*S*) and low *D* cells as ‘Receivers’ (*R*), the hexagon (H) and anti-hexagon (A) solutions are defined as (Δ*N*(*N_S_* – *N_R_*) < 0, Δ*D*(*D_S_* – *D_R_*) > 0) and (Δ*N*(*N_S_* – *N_R_*) > 0, Δ*D*(*D_S_* – *D_R_*) < 0) respectively.

**FIG. 2:**
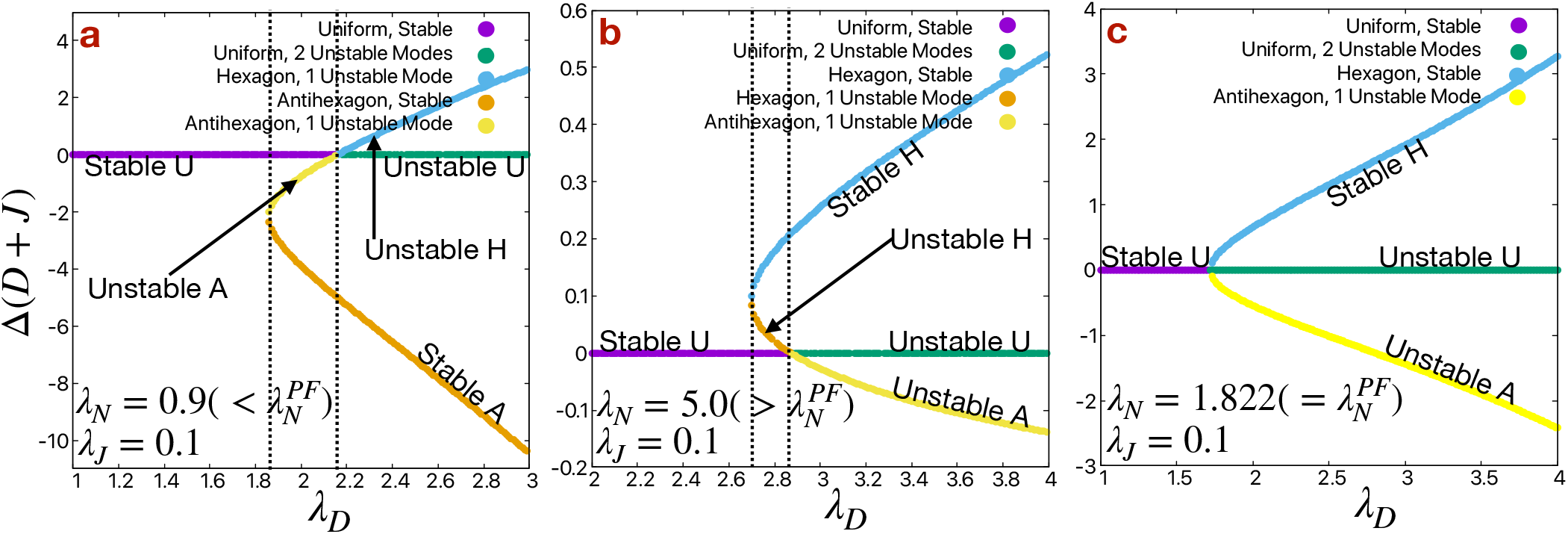
Bifurcation diagram. Bifurcation diagrams for (a) 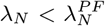, (b) 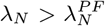 and (c) 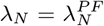 at λ_*J*_ = 0.1. All other parameters are standard.There is no practical difference between *D* + *J* versus *D* in our parameter range, so we have just refereed to the *D* pattern in the text.

Notice the proximity between the transcritical bifurcation point and a nearby saddle-node altering the stability of one or both of the hexagonal branches (Fig. 2a-b). This suggests that the system possesses a co-dimension 2 pitchfork bifurcation reflecting the coalescence of the transcritical and saddle-point bifurcations. Indeed, for a fixed value of λ_*J*_, there is always a fixed value of 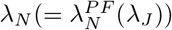 where the pitchfork bifurcation occurs; such a point is seen in Fig. 2c. At 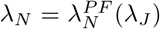, the stable uniform (U) state become unstable (with 2 unstable modes) at 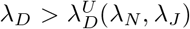 and two new states are born, a stable H and an unstable A. The interesting point is that this point appears to occur in a physically possible and experimentally observed[13] range of parameters. For all other values of λ_*N*_, the pitchfork breaks up into separate transcritical and a saddle-node bifurcations, as already discussed. For detailed discussion of this diagram for case of Notch-Delta system see[17].

Given the above, we can determine the stable states as a function of the two parameters, λ_*N*_ and λ_*D*_. The overall diagrams are presented in Fig. 3a-e for different values of λ_*J*_. The representative hexagon and antihexagon patterns are shown in the inset of Fig. 3c. For 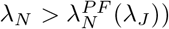 and 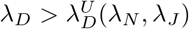, the only solutions that survive are stable hexagon (H), which is accord with the general biological finding of the absence of antihexagon phases in nature. Also, 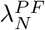 decreases with the increase in λ_*J*_, which broaden the possibility of getting hexagon phases at smaller values of λ_*N*_. On the other hand, for 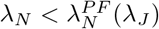 especially at higher values of λ_*J*_, there is only a small region of parameter space for which the antihexagon (A) phases are stable; this help ensure the low likelihood of antihexagon (A) phases in a biological environment with insufficient parameter control.

**FIG. 3:**
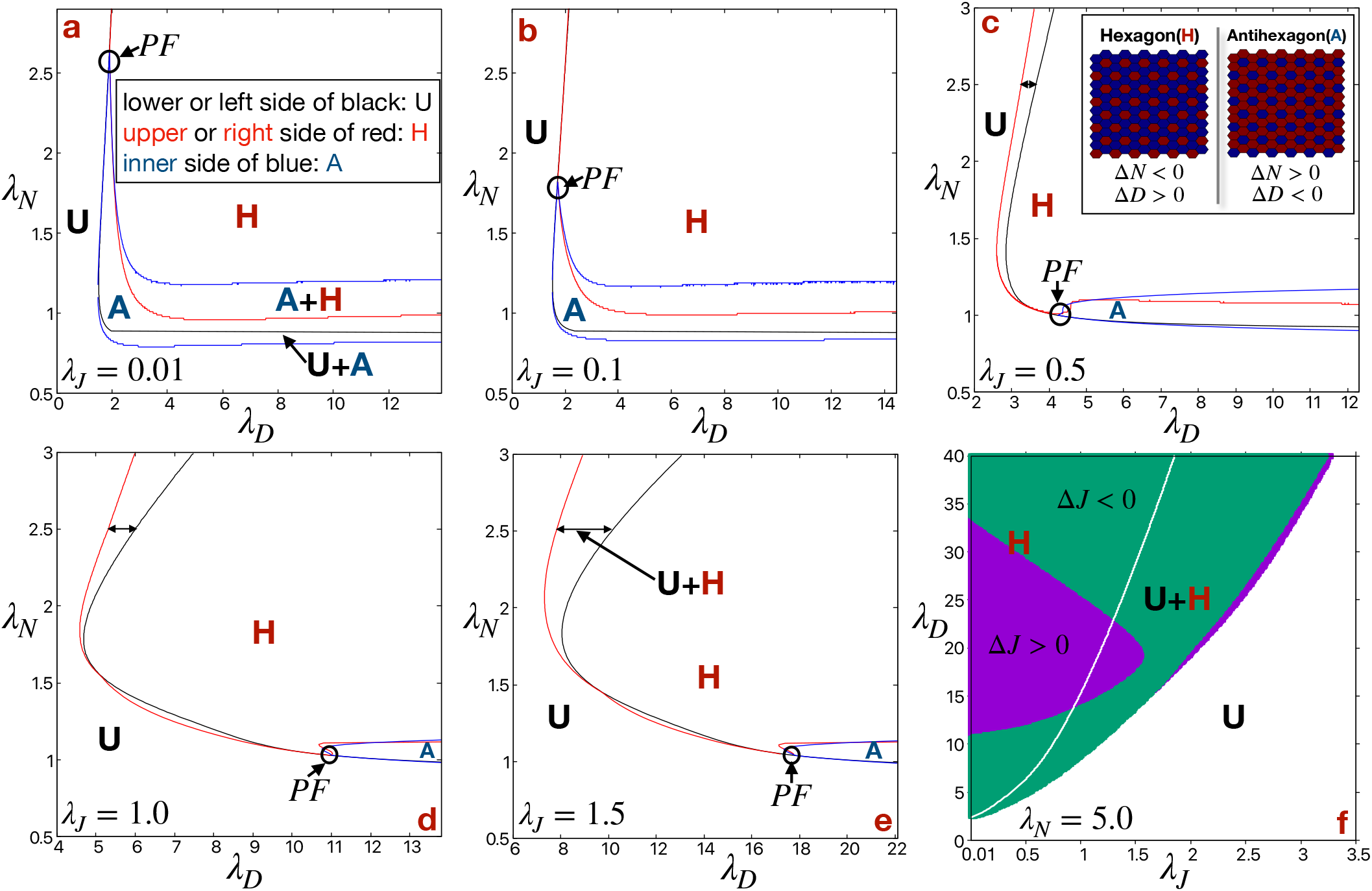
Phase diagrams. Phase diagrams in λ_*N*_ – λ_*D*_ plane consisting the regions of stable uniform (U: Δ*N*(*N_S_* – *N_R_*) = Δ*D*(*D_S_* – *D_R_*) = 0), hexagon (H: Δ*N* < 0, Δ*D* > 0), antihexagon (A: Δ*N* > 0, Δ*D* < 0) phases and different bistable regions (U+A, U+H, A+H) for (a) λ_*J*_ = 0.01, (b) λ_*J*_ = 0.1, (c) λ_*J*_ = 0.5, (d) λ_*J*_ = 1.0 and (e) λ_*J*_ = 1.5. In each phase diagram, lower or left side of black lines represents the region of stable uniform (U) phases, upper or right side of red lines represents the region of stable hexagon (H) phases, inner side of blue lines represent the region of stable antihexagon (A) phases and the point indicating by the small black circles, where U, A and H meets represent the point of pitchfork bifurcation. Inset in (c) represent the hexagon (H) and antihexagon (A) patterns of Delta (*D*) on a hexagonal lattice. (f) Phase diagram in λ_*D*_ – λ_*J*_ plane for λ_*N*_ = 5.0. The white and colored (green: *J_S_* < *J_R_* (Δ*J* = *J_S_* – *J_R_* < 0) and purple: *J_S_* > *J_R_* (Δ*J* = *J_S_* – *J_R_* > 0)) regions represent uniform (U) and hexagon (H) phases respectively. The white line represents the boundary of U region. All other parameters are standard.

Importantly, the range of parameter space where the stable uniform and stable hexagon phases coexist widens significantly as λ_*J*_ increases (Fig. 3f). Later, We will discuss the implication of this bistable region for the issue of how it could be possible to create perfectly ordered patterns.

In general, for stable patterns the *N* and *D* values are anti-correlated in a cell, whereas *N* and *I* are correlated; to wit, the cells with high *D* (labeled as Sender, *S*) will have low *N* and *I* (Fig. S2 in SI). These specific correlation ensures the fate of a cell in development, e.g., in case of development of inner ear hair cells (Senders) express high *D* and the surrounding supporting cells (Receivers) express high *N*. But *J* can be correlated or anti-correlated with *D* depending on the parameter space (in the purple region of Fig. 3f *D* and *J* are correlated, whereas in the green region they are anti-correlated). Then the immediate question arises: does this non-specific correlation between *D* and *J* affect the specification of cells’ fate? The answer appears to be no; the very much smaller difference in *J* between the Sender (*S*) and Receiver (*R*) cells with respect to the similar difference in *D* (Δ*J*(*J_S_* – *J_R_*)/Δ*D*(*D_S_* – *D_R_*) ~ 10^−2^) across our entire parameter range, ensures the lesser importance of *J* with respect to *D* for the specification of a cells’ fate.

### B. Defects in pattern formation

In the previous section, we found the regions of parameter space in which hexagon phases are stable and ordered. Considering the perfectly ordered states as a final pattern allowed us to simplify the problem to a reduced set (12) of ODEs instead of solving 4*L*^2^ ODEs on a hexagonal lattice of size *L* (total number of cells *L*^2^). The question then arises as to how these patterns can be generated in case of a realistic noisy biological environment. Starting from a uniform solution (no pattern) with small fluctuations (noise) on a hexagonal lattice of size *L* in the parameter space (λ_*N*_ = 5.0, λ_*D*_ = 10.0, λ_*J*_ = 0.5, all other parameters are standard), where the uniform state is linearly unstable, the final stable patterns are ordered for *L* = 6 (Fig. 4a) and disordered for *L* = 50 (Fig. 4b). The patterns of all the fields *N*, *D*, *J* and *I* are shown in Fig. S1 in SI. For large system size *L* = 50, the patterns are disordered with many domain boundaries formed between the hexagon patterns which nucleate and spread on different sublatices. The time evolution of *D* for all the cells is shown in Fig. 4c, Video S1 (for *L* = 6) and Video S2 (for *L* = 50).

**FIG. 4:**
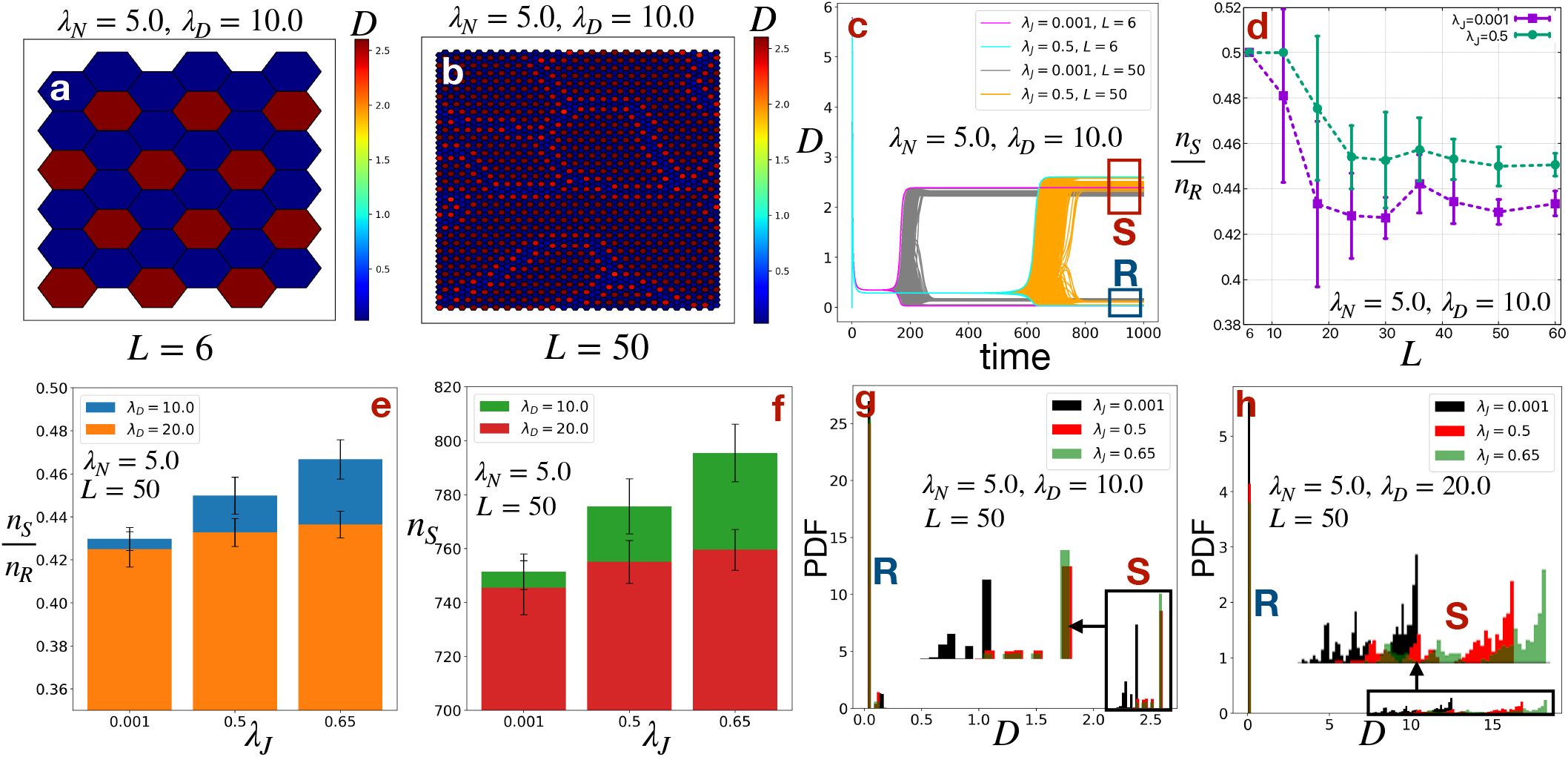
Defects in patterns. Steady state patterns of Delta (*D*) at λ_*N*_ = 5.0, λ_*D*_ = 10.0, λ_*J*_ = 0.5 for two different system sizes (a) *L* = 6 (defect-free) and (b) *L* = 50 (defective). (c) Dynamics of Delta (*D*) for all the cells in a hexagonal lattice of size *L* = 6 and *L* = 50 at λ_*N*_ = 5.0, λ_*D*_ = 10.0, λ_*J*_ = 0.001 and 0.5. The branches enclosed in the red and blue rectangles represent the Sender (*S*: high *D*) and Receiver (*R*: low *D*) cells respectively. (d) The ratio of number of Sender (S) and Receiver (R) cells in steady states 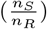 as a function of system size *L* at λ_*N*_ = 5.0, λ_*D*_ = 10.0, λ_*J*_ = 0.001 and 0.5. (e) The same ratio 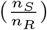 and (f) number of Sender (*S*) states (*n_S_*) for *L* = 50 at λ_*J*_ = 0.001, 0.5 and 0.65 for different values of λ_*D*_ at λ_*N*_ = 5.0. Probability density (PDF) of Delta (*D*) in steady states for system size *L* = 50 at (g) λ_*N*_ = 5.0, λ_*D*_ = 10.0 and (h) λ_*N*_ = 5.0, λ_*D*_ = 20.0. The insets in (g) and (h) show the enlarged version of the PDF of the Sender (*S*) states. All other parameters are standard.

The ratio between high *D* (*S*) and low *D* (*R*) cells 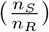 roughly estimates the degree of disorder in steady state patterns. For a perfectly ordered pattern this ratio should be close to 0.5 (for *L* = 50, *n_S_* = 825 and *n_R_* = 1675). 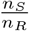 decreases gradually as *L* increases and starts to saturate for *L* > 18 (Fig. 4d). For a larger system the nucleation centers for hexagonal patterns are more plentiful, which leads to higher disorder in the patterns. But, 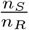 increases with the increase in λ_*J*_ across different *L* (Fig. 4d) and λ_*D*_ (Fig. 4e). The time evolution and steady state patterns of *D* for different values of λ__D__ are shown in Fig. S2 in SI. The increase in 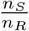 or *n_S_* (Fig. 4e) with increase in λ_*J*_ and/or decrease in λ_*D*_ indicates that we can find the maximum number of Sender cells near the stability boundary of the Uniform (U) phase (the white line in the phase diagram in Fig. 3f). Similarly, the phase diagrams in Fig. 3a-e also suggest that the *n_S_* should increase as λ_*N*_ decreases. We do observe a gradual increment in *n_S_* or 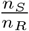 as λ_*N*_ decreases (Fig. S3a,c in SI). For very small values of λ_*N*_ = 1.5, we find a different kind of disordered pattern where two neighboring cells can have high *D* (Fig. S3a in SI, Video S3), which leads to *n_S_* > 825 or 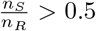. This situation can also be remediated by increasing λ_*J*_. That is, higher λ_*J*_, at this smaller value of λ_*N*_ = 1.5, decreases the possibility of getting two neighboring high *D* cells and hence leads towards more ordered patterns (Fig. S3b,d in SI).

We also observed a wide distribution of *D* values among the high *D*, Sender (*S*) cells (Fig. 4g-h). The long tail in the distributions arises from the cells at the hexagonal domain boundaries of the disordered patterns. Conversely, most of the cells at the core of the hexagonal domains express similar values of *D* to the values of *D* for perfectly ordered patterns, as we found for *L* = 6 (Fig. 4c). In general, we need additional strategies, beyond shifting the system such that parameters lie closer to the boundary of stable uniform solutions, in order to get defect-free patterns in a biological system, we will return to this below. But, an increment in the number of high *D* cells with increase in λ_*J*_ near the boundary of the stable uniform regions of the phase spaces, may have other implications in biological systems. For example, in case of collective migration larger number of high *D* cells can increase the overall invasiveness (highly invasive or leader cells express high *D*).[21, 22]

Moreover, the time to reach the steady state (differentiation time) increases as λ_*J*_ increases (Fig. 4c). This is again because higher λ_*J*_ shifts the system towards stability of the Uniform state, which then increases the differentiation time. This slow differentiation time can accommodate other slow processes of error correction in the pattern formation such as a large delay in protein production.[23]

### C. Dose-dependent role of Jagged

Depending on the production rate of Jagged (λ_*J*_) the cells can attain different fates. Starting from a lateral inhibition pattern of high *D* (low *I*, low *N*) Sender (*S*) and low *D* (high *I*, high *N*) Receiver (*R*) cells at small values of λ_*J*_, a hybrid (*S/R*) state with intermediate values of *I* [14, 15] appears as λ_*J*_ increases (Fig. 5a). Note that at low λ_*J*_, where the Notch-Delta signaling dominates, Jagged acts synergistically with Delta to refine the lateral inhibition pattern of Sender and Receiver cells. The addition of Jagged to Delta in binding the common resource of Notch receptors leads to the greater activation of NICD and hence stronger suppression of Delta in the neighboring Receiver cells. We quantify this by calculating the difference in Delta (Δ*D*) between the Sender and Receiver cells (*D_S_* – *D_R_*) as a function of λ_*J*_, considering the perfectly ordered patterns where at steady state all the Sender and Receiver cells attain specific *D_S_* and *D_R_* values respectively. We observe a non-monotonic dependence of Δ*D* as a function of λ_*J*_ across different values of λ_*D*_ (Fig. 5b). Up to a certain value of λ_*J*_, Δ*D* increases gradually and reaches a maximum; with the further increase in λ_*J*_ the strength of Notch-Delta and Notch-Jagged signaling becomes comparable, the system enters into a region of bistability where both the uniform (U) and hexagon (H) phase are stable, and eventually Δ*D* starts to decrease gradually. Both *D_R_* and *D_S_* increases as λ_*J*_ increases up to the critical value of λ_*J*_ (Fig. S4 in SI), but the increment in *D_S_* always much higher than the increment in *D_R_*. At a fixed value of λ_*J*_, the competition between *D* and *J*, and thus the value of Δ*D* can be enhanced by either increasing λ_*D*_ at fixed λ_*N*_ (Fig. 5b) or decreasing λ_*N*_ at fixed λ_*D*_ (Fig. S4d-e in SI). At very high values of λ_*J*_, lateral induction dominates and the pattern becomes uniform consisting entirely of hybrid *S/R* cells. Hybrid *S/R* states have an critical role in promoting collective migration in wound healing [24] and cancer metastasis.[15] This dose-dependent role of Jagged is shown in the schematic diagram Fig. 5c.

**FIG. 5:**
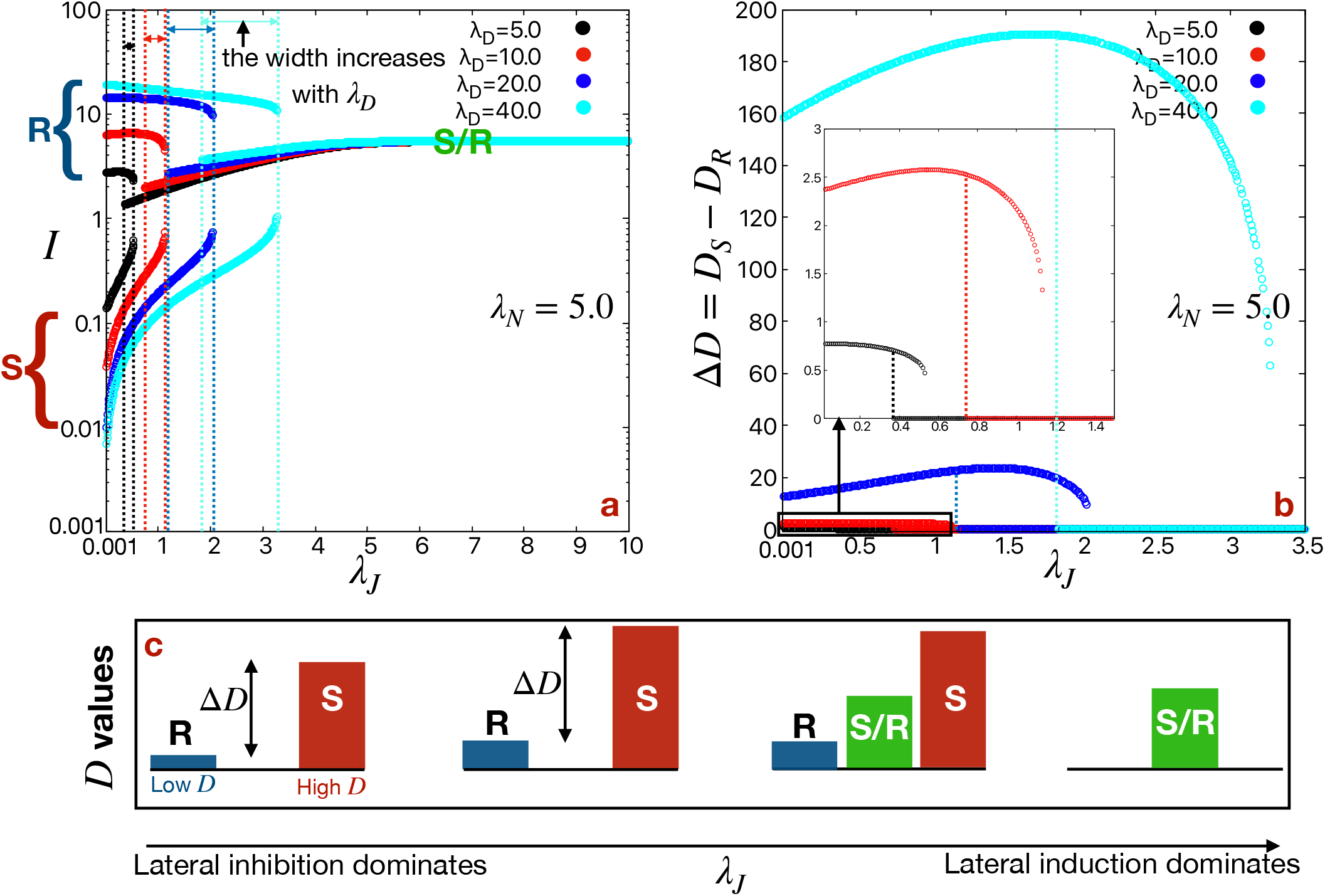
Dose-dependent role of Jagged (*J*). (a) NICD (I) as a function of λ_*J*_ showing the Sender (*S*: high *D*), Receiver (*R*: low *D*) and hybrid *S/R* (intermediate *D*) state branches. (b) The difference in Delta (Δ*D*) between the *S* and *R* cells (*D_S_* – *D_R_*) as a function of λ_*J*_. The inset shows an enlarged version of the rectangular region at very small values of Δ*D*. All other parameters are standard. (c) Schematic representation of *R*, *S* and *S/R* states and their Delta (D) values as a function of λ_*J*_.

This synergistic role of Jagged with Delta enabling the robust lateral inhibition pattern of high *D* hair cells surrounded by low *D* supporting cells has been suggested experimentally in the hair cell differentiation phase of chick inner ear development.[7] This idea has also been proposed in a model of angiogenesis to enable the robust patterning of high *D*, Tip and low *D*, Stalk cells.[25, 26]

### D. Effect of cis-inhibition and trans-activation strength

Although cis-inhibition (*k_c_*) does not directly contribute to the production of NICD signal, it affects the patterns by altering the Notch, Delta and Jagged expressions. It has been shown that *k_c_* increases the robustness of lateral inhibition patterns by inactivating Notch in Sender cells. At first, we draw a phase diagram in λ__D__ – *k_c_* plane for a smaller value of λ_*J*_ = 0.1 (Fig. S5a in SI). We find a new kind of surprising stable state with *N* and *D* are correlated for very smaller values of *k_c_* (cyan and orange colored region in phase diagram in Fig. S6a). As opposed to the usual anti-correlation of *N* and *D* values in a cell, here both Δ*N*(*N_S_* – *N_R_*) > 0 and Δ*D*(*D_S_* – *D_R_*) > 0. We refer these solutions as High-High (Hi-Hi), since both the *D* and *N* are higher in Sender cells compared to those in Receiver cells. As discussed in,[17] these kind of states have not observed experimentally to date, presumably because the parameters for which these solutions exist are not typically found in a developmental process.[13] Apart from that, at higher λ_*J*_ these Hi-Hi states do not exist even at smaller values of *kc* as the patterns become uniform (Fig. 6a).

**FIG. 6:**
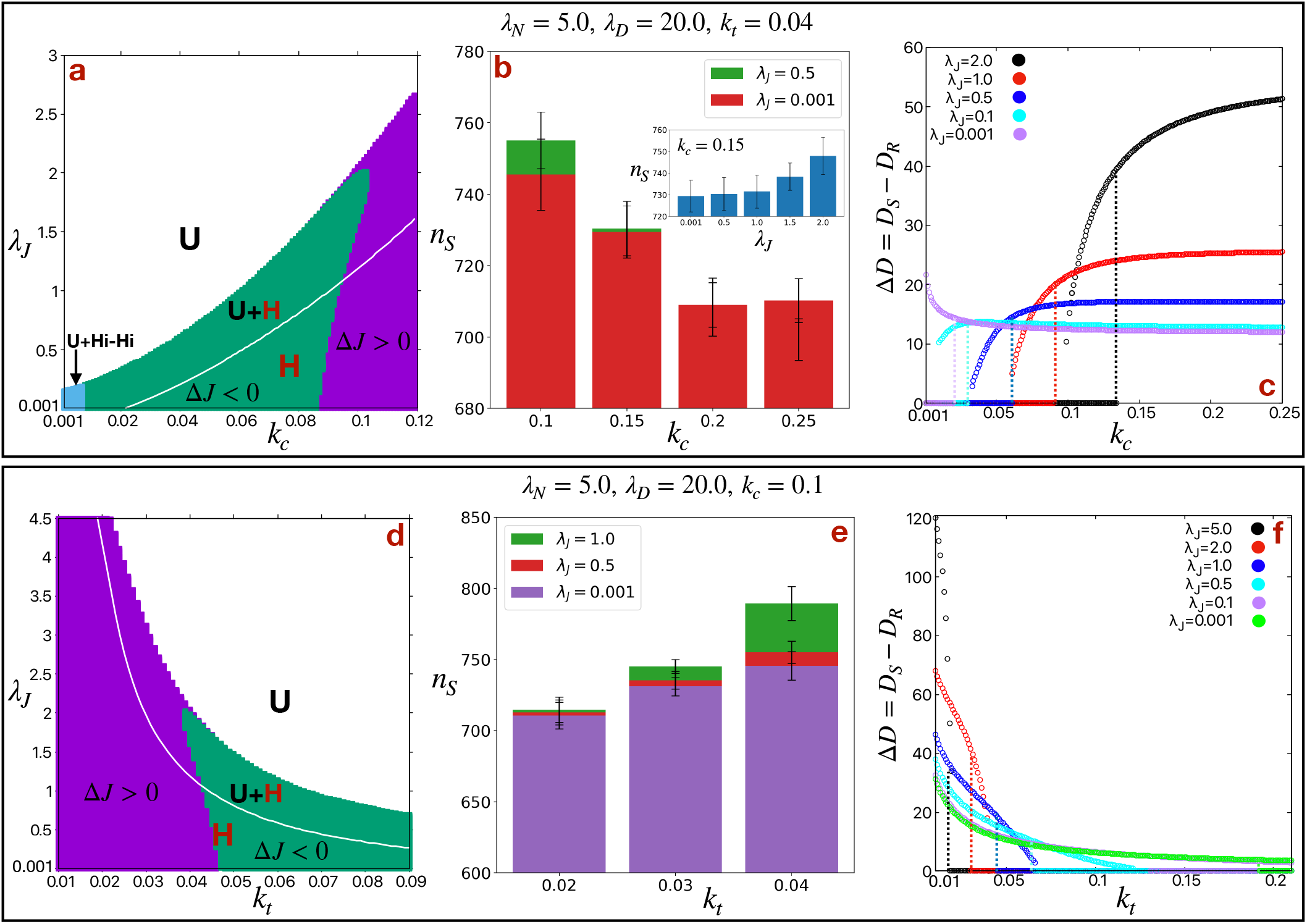
Role of cis-inhibition (*k_c_*) and trans-activation (*k_t_*). (a) Phase diagrams in λ_*J*_ – *k_c_* plane for λ_*N*_ = 5.0, λ_*D*_ = 20.0 and *k_t_* = 0.04. The white and colored (green: Δ*J*(*J_S_* ‒ *J_R_*) < 0) and purple: (Δ*J*(*J_S_* – *J_R_*) > 0)) regions represent uniform (U: Δ*N*(*N_S_* – *N_R_*) = Δ*D*(*D_S_* – *D_R_*) = 0) and hexagon (H: Δ*N* < 0, Δ*D* > 0) phases respectively. The cyan colored region represents the bistability of uniform (U) and High-High (Hi-Hi: Δ*N* > 0, Δ*D* > 0) phases. The white line represents the boundary of U region. (b) The number of Sender (S) states (*n_S_*) and (c) the difference in Delta (Δ*D*) between the Sender (*S*) and Receiver (*R*) states (*D_S_* – *D_R_*) as a function of *k_c_* for different values of λ__J__ at λ_*N*_ = 5.0, λ_*D*_ = 20.0, *k_t_* = 0.04 for a hexagonal lattice of size *L* = 50. The inset in (b) shows the number of Sender (*S*) cells (*n_S_*) as a function of λ_*J*_ at a fixed value of *k_c_* = 0.15. The similar (d) phase diagram in λ_*J*_ – *k_t_* plane, (e) number of Sender (*S*) cells (*n_S_*) as a function of *k_t_* and (f) difference in Delta (Δ*D*) between the Sender (*S*) and Receiver (*R*) cells (*D_S_* – *D_R_*) as a function of *k_t_* for λ_*N*_ = 5.0, λ_*D*_ = 20.0, *k_c_* = 0.1. All other parameters are standard.

As *k_c_* increases, we move further away from the boundary of the stable uniform phases (white line in the phase diagram in Fig. 6a), which decrease the number of Sender cells (*n_S_*) by creating more disordered states (Fig. 6b). This can be again remediated by increasing λ_*J*_ as shown in Fig. 6b. Also as *k_c_* increases, higher value of λ_*J*_ is needed to get stable hexagon patterns (Fig. 6a). The higher values of λ_*J*_ allow higher values of Δ*D* (at fixed *k_c_*), and hence increases the robustness of the patterning. In general, Δ*D* increases with increase in *k_c_* up to a certain value of *k_c_* and then starts to saturate (Fig. 6c). Surprisingly, ΔD decreases as *k_c_* increases for very smaller value of *λ_J_* = 10^−3^. Actually, this behavior depends on the relative availability of Notch and Delta. Thus, the dependence of Δ*D* on *k_c_* can be switched by either increasing λ_*D*_ at fixed λ_*N*_ (Fig. S5b in SI) or decreasing λ_*N*_ at fixed λ_*D*_ (Fig. S5c in SI).

As opposed to *k_c_*, the trans-activation strength (*k_t_*) interacts with λ_*J*_ in an opposite manner. Smaller values of *k_t_* broaden the range of λ_*J*_ for which hexagon phases are stable (Fig. 6d). Furthermore, *n_S_* increases and Δ*D* decreases as *k_t_* increases. In short, in presence of higher λ_*J*_, higher values of *k_c_* and/or lower values of *k_t_* increase the robustness (higher values of Δ*D*) of the patterns. We note that it has been observed experimentally that the lateral inhibition patterning in the zebrafish notochord is promoted when ligand-receptor interactions are stronger within the same cell (*k_c_*) than in neighboring cells (*k_t_*).[27]

### E. Defect-free patterns, revisited

As discussed earlier, pattern arising from the uniform states with small noise are in general disordered, with many domains of hexagon patterns. The hexagon patterns randomly nucleate at different sublattices and spread over time in a disordered manner. One way to avoid such disorder can be to find a parameter set for which a local perturbation which nucleates the pattern would spread over time in a ordered manner to create a perfectly ordered pattern.[28] This can happen in the bistable region, where both the uniform and hexagon phases are stable (Fig. 3).

Fig. 7 shows the spatiotemporal patterns of *D* on a hexagonal lattice starting from a hexagonal seed in the center of the lattice, for different parameters. For the parameter space (λ_*N*_ = 5.0, λ_*D*_ = 10.0, λ_*J*_ = 0.5, all other parameters are standard) where only the hexagon phase is stable, as expected the pattern nucleates at different sublattices and spreads over time in a disordered manner (Fig. 7a, Video S4). But, for the parameter set (λ_*N*_ = 5.0, λ_*D*_ = 10.0, λ_*J*_ = 0.9, all other parameters are standard) chosen from the region of bistability, the initial hexagonal seed spreads over time in a ordered manner and thereby creates perfectly order pattern (Fig. 7b, Video S5).

**FIG. 7:**
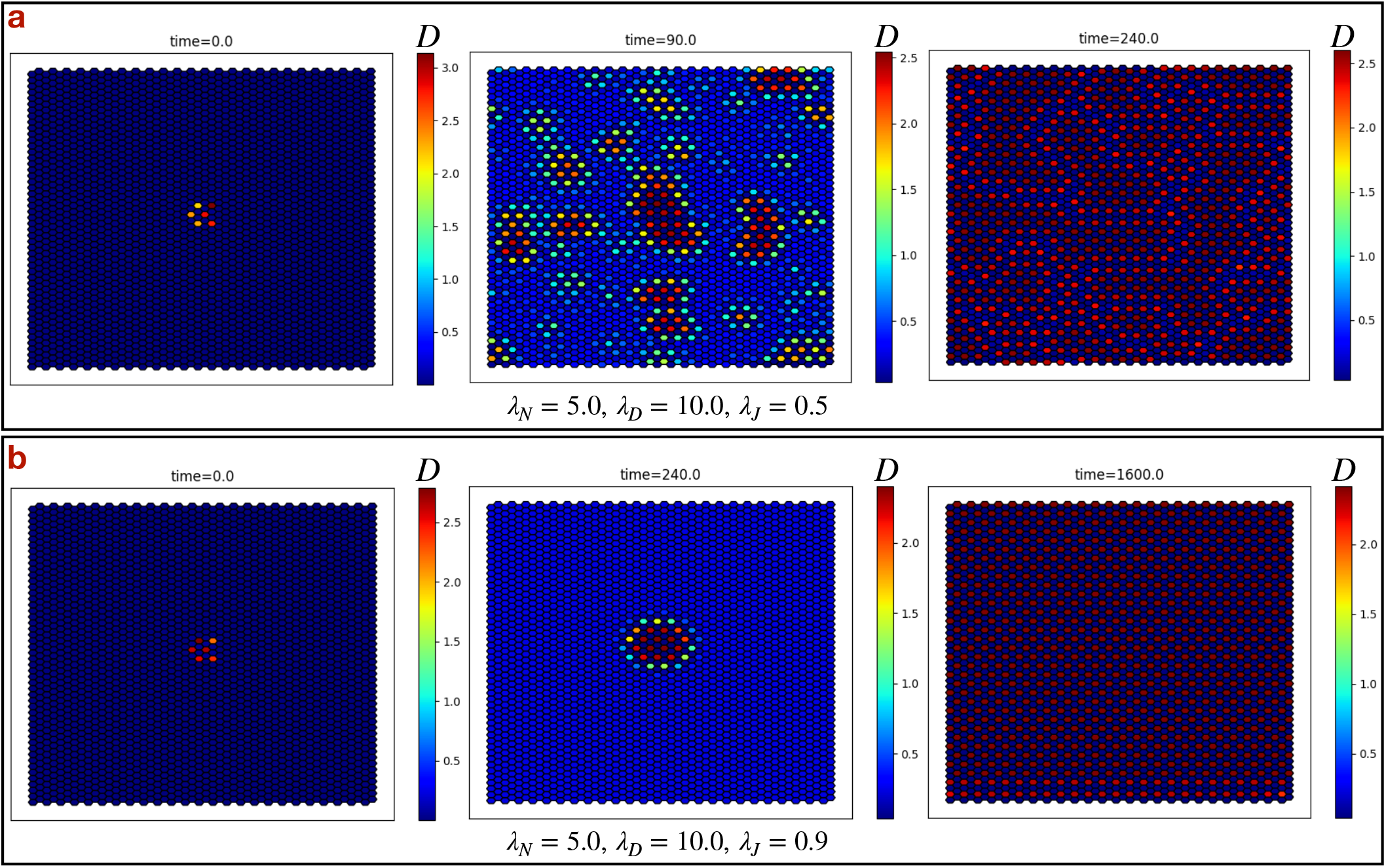
Spatiotemporal patterns of Delta (*D*) starting from a hexagonal seed. (a) The parameters (λ_*N*_ = 5.0, λ_*D*_ = 10.0, λ_*J*_ = 0.5) for which only the Hexagon (H) phase is stable the steady state patterns are imperfect, disordered. (b) The parameters (λ_*N*_ = 5.0, λ_*D*_ = 10.0, λ_*J*_ = 0.9) for which both the Hexagon (H) and Uniform (U) phases are stable (bisatble region) the steady state patterns are defect-free, ordered. All other parameters are standard.

In generic Notch-Delta system without Jagged, the parameter range exhibiting this bistability is very narrow (Fig. 3a). But, including finite λ_*J*_ widens the range significantly (Fig. 3f). We can also investigate the effect of different parameters on the width of the region of bistability by computing the phase diagrams in the λ_*D*_ – λ_*J*_ plane (Fig. S6 in SI). We observe that region of bistability in the phase space increases, especially at higher values of λ_*D*_ and λ_*J*_, as λ_*N*_ increases and/or *k_c_* decreases and/or *k_t_* increases. Thus, Jagged helps to widen the the bistable region, which can help to get ordered hexagon patterns in biological system with weaker control of operating parameters.

## IV. CONCLUSIONS

In this paper, we explored pattern formation in the Notch-Delta-Jagged signaling on multicellular system. Assuming hexagonal symmetry of a cellular lattice (as being close to biological tissue) helps us to compute the phase space by reducing the problem of ordered patterns to a set of 12 coupled ODEs. Throughout the paper, we focus on the role of Jagged on the accuracy and robustness of the pure Notch-Delta pattern. We observe that Jagged decreases the possibility of obtaining nonphysical antihexagon states by shrinking the parameter range for which antihexagon solutions are stable. Higher production of Jagged (λ_*J*_) also ensures the absence of experimentally unseen Hi-Hi states (where both Notch and Delta are high in Sender Cells) at small values of the cis-inhibition rate (*k_c_*).

In general, starting from a uniform state with small fluctuations, incommensurate hexagon patterns emerges on different sublattices and the pattern spreads in a disordered manner; the final lateral induction pattern contains many domain boundaries between the hexagon structures. We quantified this disorder by calculating the number of Sender cells (*n_S_*) in the lattice. For a ordered lattice of size *L* (total number of cells = *L*^2^), *n_S_* should be around *L*/3. Table I summarizes the effect of different parameters on *n_S_*. At a fixed value of the other parameters, *n_S_* decreases with the production rate of Notch (λ_*N*_) and/or production rate of Delta (λ_*D*_) and/or cis-inhibition rate (*k_c_*) increases, but *n_S_* increases as trans-activation rate (*k_t_*) increases. In all cases, *n_S_* increases as production rate of Jagged (λ_*J*_) increases (except for very smaller values of λ_*N*_), which leads to a more ordered pattern. As a general rule, *n_S_* is maximum near the stability boundary of the Uniform phase. Similarly, the time to reach the steady state (the differentiation time) is maximum near the boundary of the Uniform phase, which can be reached by changing the parameters as shown in Table I. The slow differentiation may allow for other mechanisms to resolve defects in the pattern.

**TABLE I:**
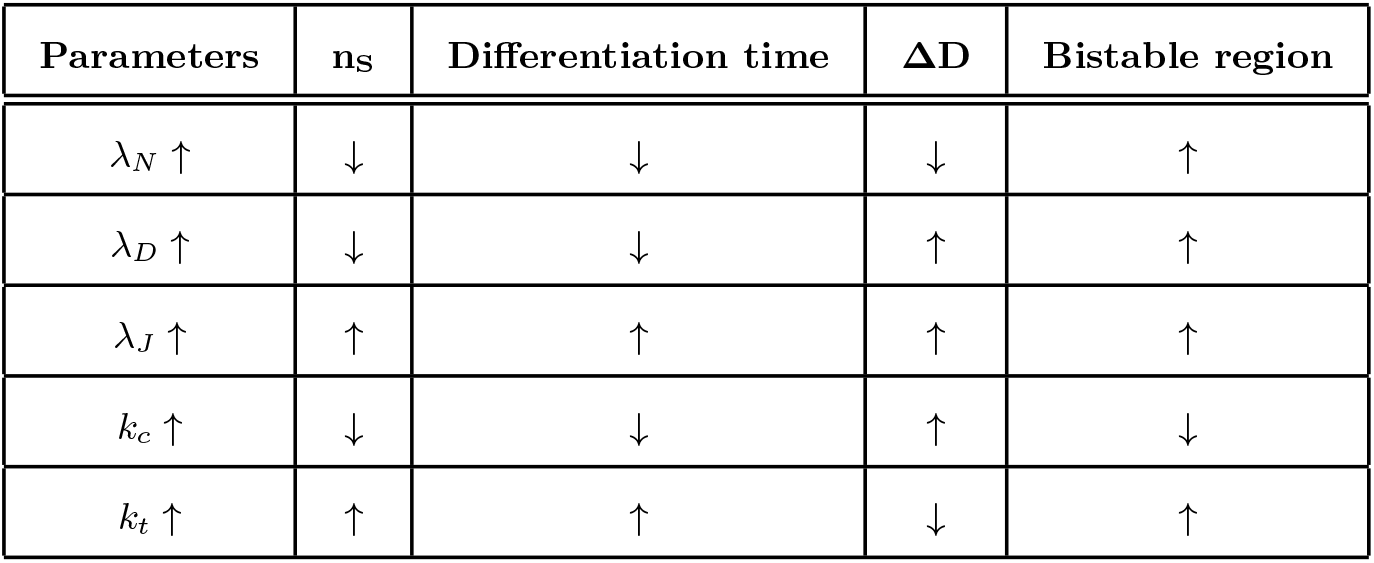
Effects of different parameters on stable hexagon pattern formation.

| Parameters | $n_S$ | Differentiation time | $\Delta D$ | Bistable region |
| --- | --- | --- | --- | --- |
| $\lambda_N \uparrow$ | ↓ | ↓ | ↓ | ↑ |
| $\lambda_D \uparrow$ | ↓ | ↓ | ↑ | ↑ |
| $\lambda_J \uparrow$ | ↑ | ↑ | ↑ | ↑ |
| $k_c \uparrow$ | ↓ | ↓ | ↑ | ↓ |
| $k_t \uparrow$ | ↑ | ↑ | ↓ | ↑ |

We quantified the robustness of a pattern by calculating the difference in Delta (Δ*D*) between the Sender and Receiver cells (*D_S_* – *D_R_*). Δ*D* increases as λ_*J*_ increases up to the point where Notch-Delta signaling no longer dominates over Notch-Jagged signaling. The competition between Delta and Jagged over binding with Notch increases the Delta in Sender cells compared to the Receiver cells. Δ*D* also increases as λ_*D*_, *k_c_* increases and λ_*N*_, *k_t_* decreases (Table I).

As listed in Table I, at sufficiently higher values of λ_*N*_, λ_*D*_, λ_*J*_, *k_t_* and smaller value of *k_c_*, a large bistable region consisting of uniform and hexagon phases allows the emergence of defect-free patterns. Without Jagged-mediated signaling, this bistable region is very narrow. It would be difficult for a biological system to reliably adjust the parameters to lie within the bistable region in the case of small λ_*J*_. In the pure Notch-Delta system without Jagged, many strategies has been proposed throughout the literature to get biologically relevant defect-free patterns, by adjusting the time delays,[23, 29] the noise[30, 31] in the network, coupling a parameter to an initiation wave,[17] or coupling to different properties of cells with a core Notch-Delta circuit such as apoptosis,[32] cell cycle,[33] adhesion,[34, 35] cell mechanics[36–38] etc. The mechanisms mentioned above, along with the Jagged-mediated broadening of bistable region, should lead to interesting future studies in the Notch-induced pattern formation problem.

## Author Contributions

M.M. and H.L. designed the study. M.M. developed the code, performed simulations and analyze the data. M.M. and H.L. prepared, reviewed and finalized the manuscript.

## Conflicts of interest

There are no conflicts to declare.

## Acknowledgements

This work was supported by National Science Foundation by sponsoring the Center for Theoretical Biological Physics – award PHY-2019745 and also by award PHY-1605817.

## Supplementary Information

### The description of the video files

Video S1: Spatiotemporal patterns of Delta (*D*) starting from uniform initial condition with small fluctuations for λ_*N*_ = 5.0, λ_*D*_ = 10.0, λ_*J*_ = 0.5 and *L* = 6. All other parameters are standard.

Video S2: Spatiotemporal patterns of Delta (*D*) starting from uniform initial condition with small fluctuations for λ_*N*_ = 5.0, λ_*D*_ = 10.0, λ_*J*_ = 0.5 and *L* = 50. All other parameters are standard.

Video S3: Spatiotemporal patterns of Delta (*D*) starting from uniform initial condition with small fluctuations for λ_*N*_ = 1.5, λ_*D*_ = 10.0 and λ_*J*_ = 0.1. All other parameters are standard.

Video S4: Spatiotemporal patterns of Delta (*D*) starting from a hexagonal seed for λ_*N*_ = 5.0, λ_*D*_ = 10.0 and λ_*J*_ = 0.5. All other parameters are standard.

Video S5: Spatiotemporal patterns of Delta (*D*) starting from a hexagonal seed for λ_*N*_ = 5.0, λ_*D*_ = 10.0 and λ_J_ = 0.9. All other parameters are standard.

### The reduced set of 12 ODE’s

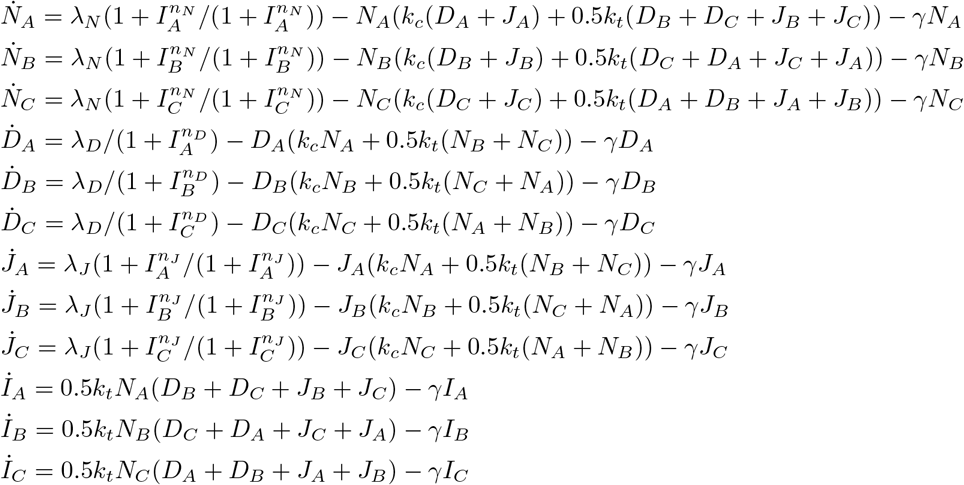

**TABLE S1:**
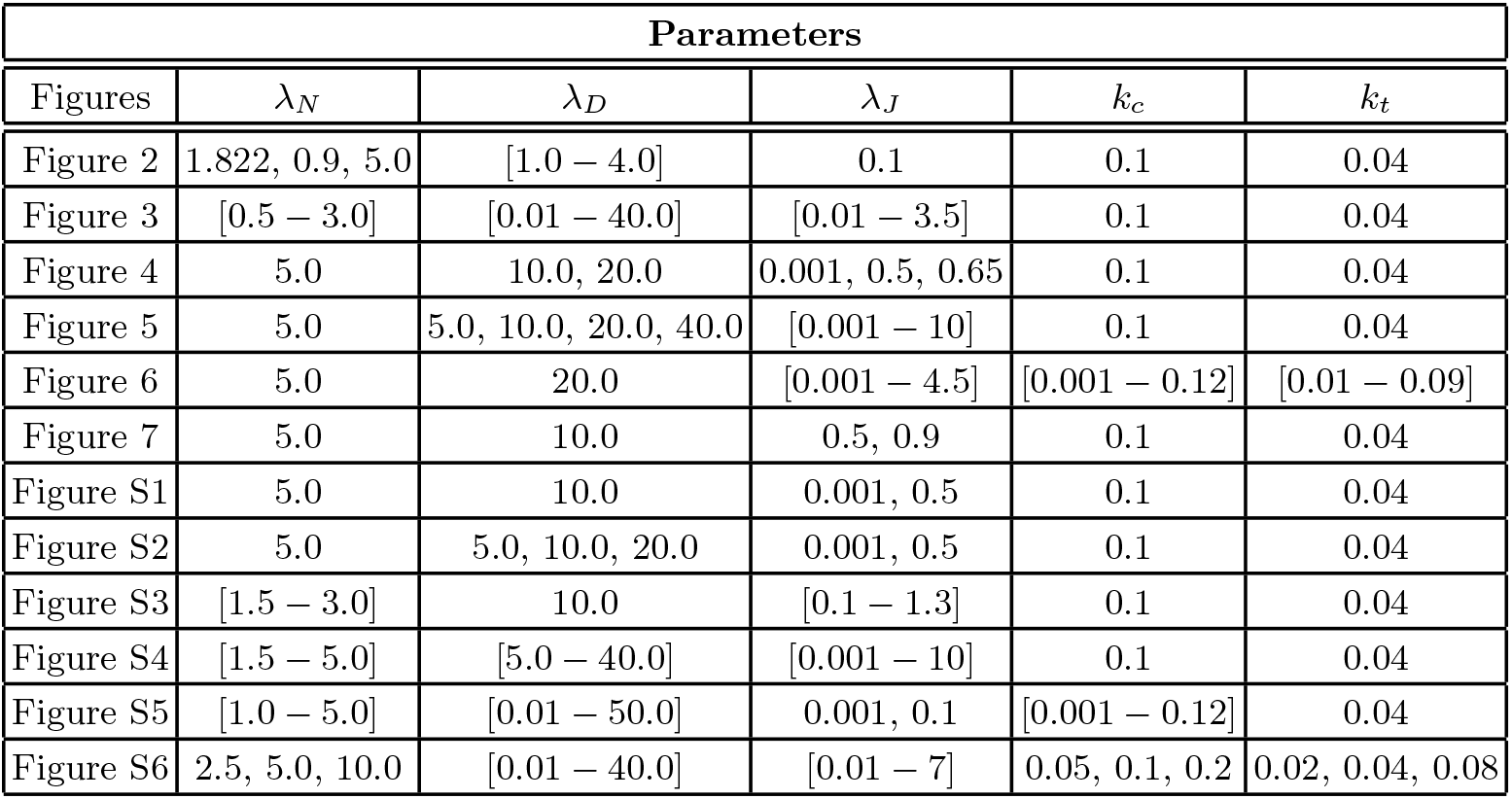
Parameters for all the Figures described in the manuscript.

**FIG. S1:**
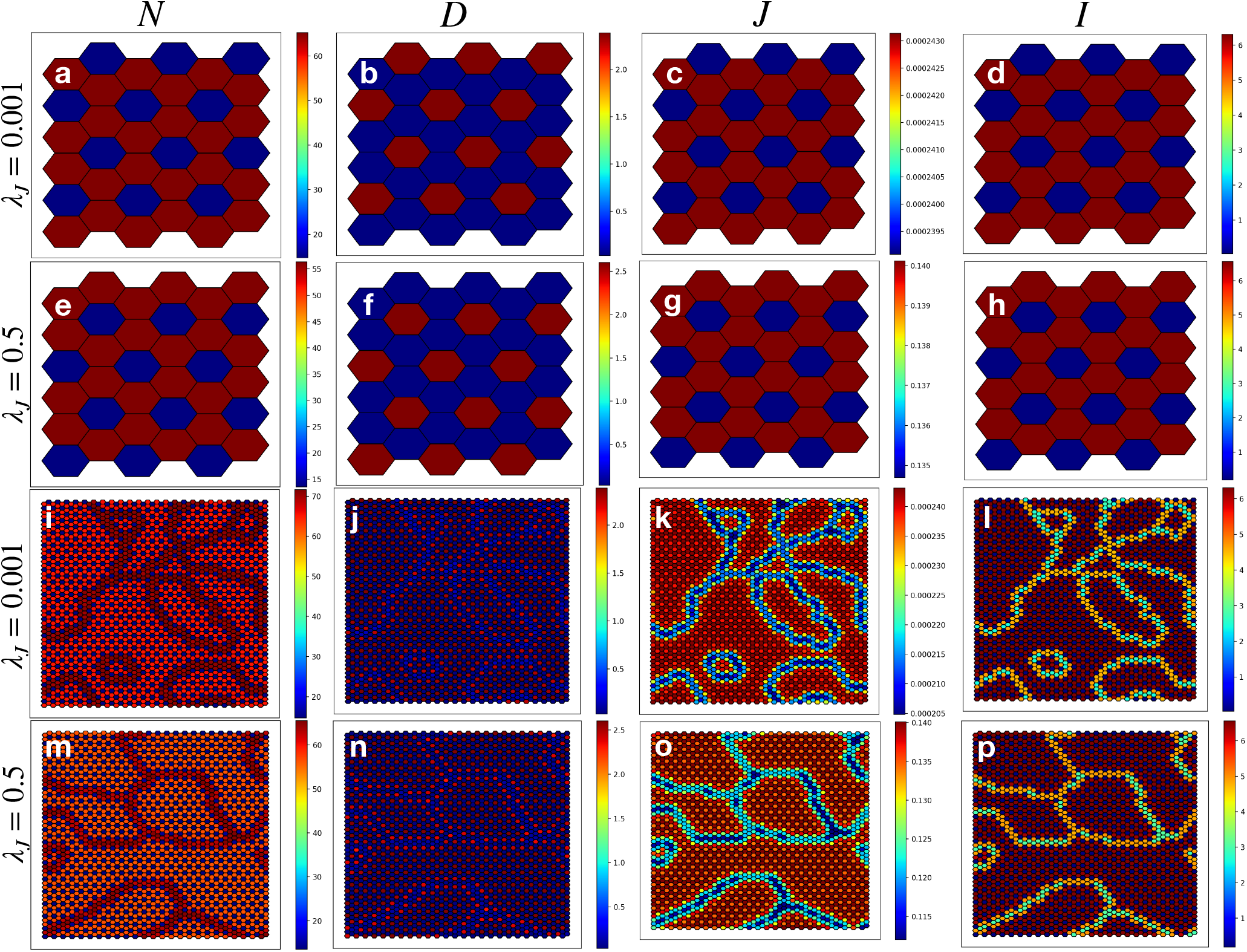
Steady state patterns of Notch (N), Delta (D), Jagged (J) and NICD (I) on a hexagonal lattice. The steady states of Notch (N), Delta (D), Jagged (J) and NICD (I) at λ_*N*_ = 5.0, λ_*D*_ = 10.0 and λ_*J*_ = 0.001 and 0.5 on a hexagonal lattice of size (a–h) *L* = 6 and (i–p) *L* = 50. All other parameters are standard.

**FIG. S2:**
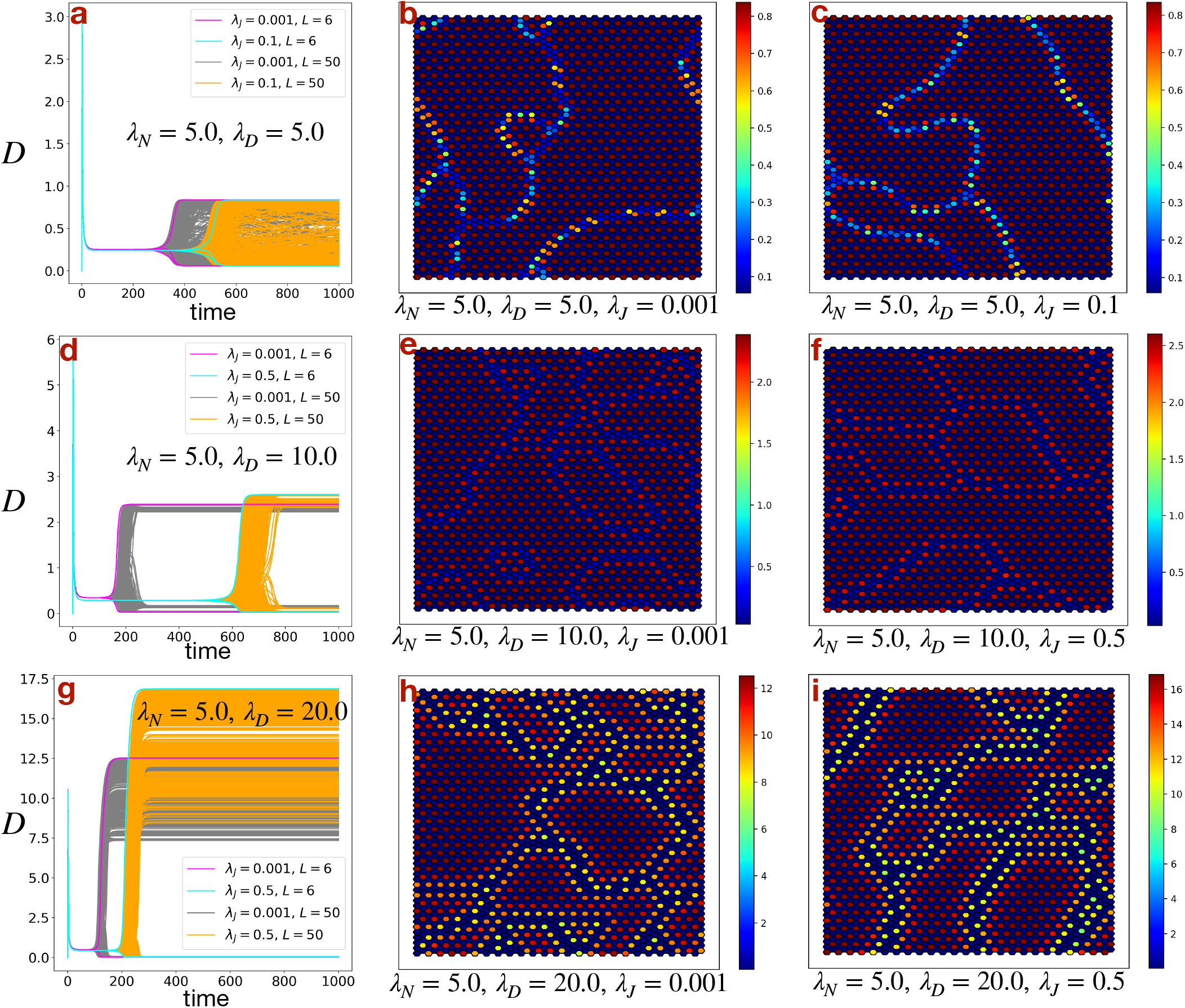
Effect of production rate of Delta (λ_*D*_). Dynamics of Delta (D) for all the cells on a hexagonal lattice of size *L* = 6 and *L* = 50 at λ_*N*_ = 5.0, λ_*J*_ = 0.001 and 0.5 (0.1 instead of 0.5 for λ_*D*_ = 5.0, because at λ_*J*_ = 0.5, λD = 5.0 and λ_*N*_ = 5.0, the steady states become uniform (U)) for (a) λ_*D*_ = 5.0, (d) λ_*D*_ = 10.0 and (g) λ_*D*_ = 20.0. (b), (c), (e), (f), (h), (i) Corresponding patterns of Delta (D) at steady states for *L* = 50 for different values of λ_*D*_ and λ_*J*_. All other parameters are standard.

**FIG. S3:**
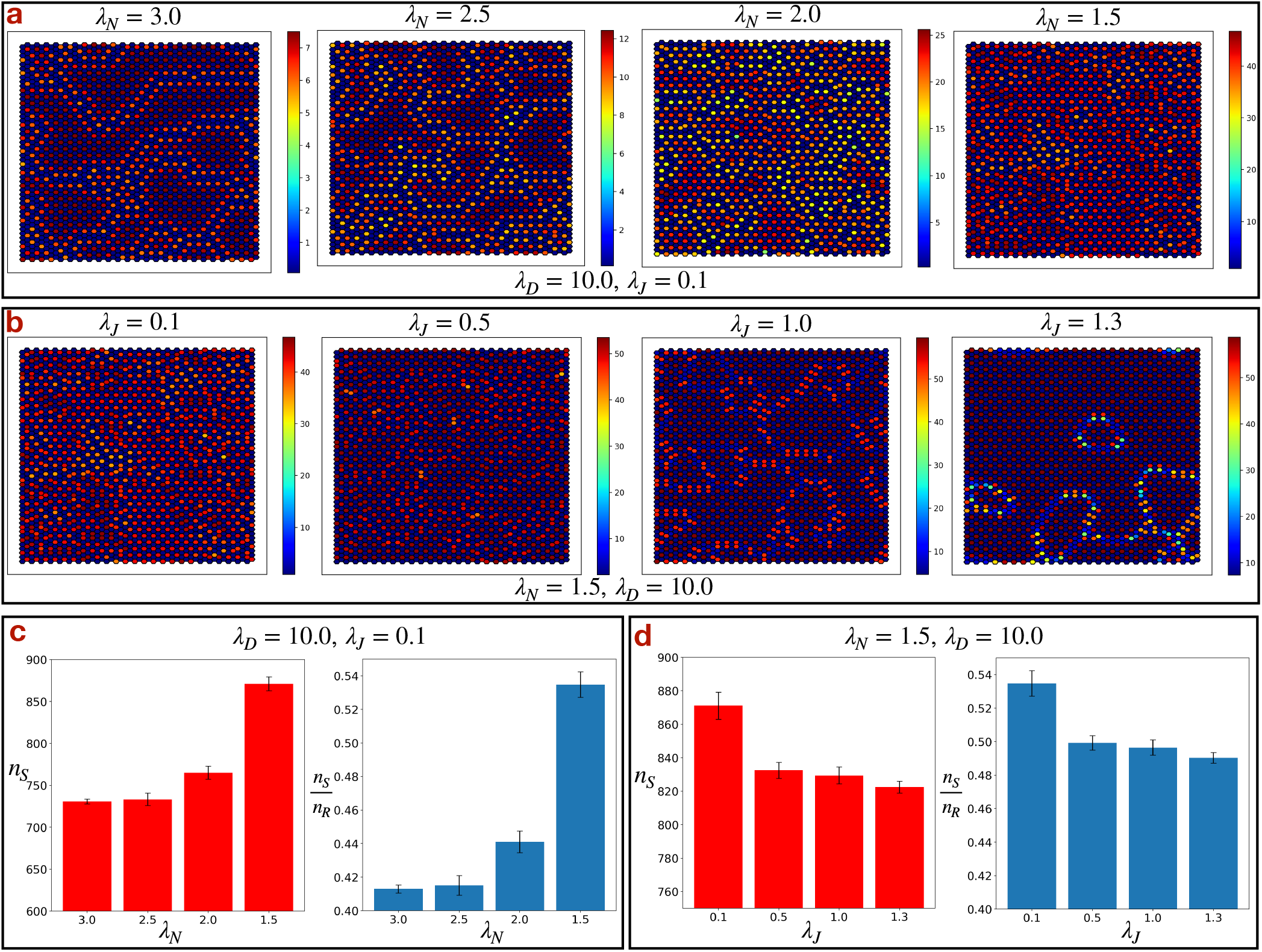
Effect of production rate of Notch (λ_*N*_). The steady state patterns of Delta (*D*) on a hexagonal lattice of size *L* = 50 (a) for varying λ_*N*_ at λ_*D*_ = 10.0, λ_*J*_ = 0.1 and (b) for varying λ_*J*_ at λ_*N*_ = 1.5, λ_*D*_ = 10.0. (c), (d) The corresponding number of Sender (S) states (*n_S_*) and ratio of number of Sender (S) and Receiver (R) states 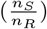. All other parameters are standard.

**FIG. S4:**
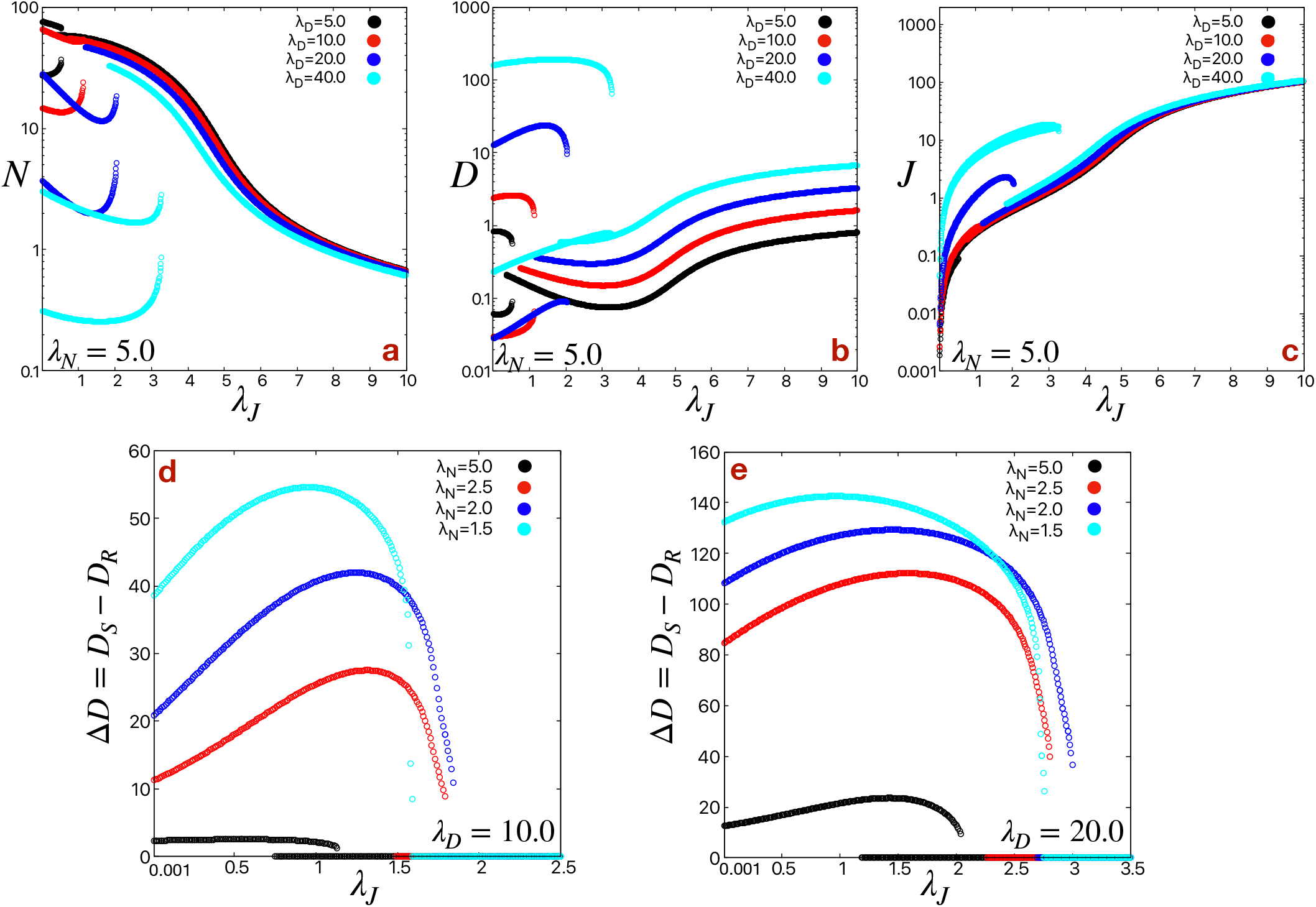
Steady state values of Notch (N), Delta (D) and Jagged (J) as a function of λ_*J*_. Steady state values of (a) Notch (N), (b) Delta (D), (c) Jagged (J) as a function of λ_*J*_ for different values of λ_*D*_ at λ_*N*_ = 5.0. The y-axes (*N*, *D* and *J*-axes) are in log-scale. The difference in Delta (Δ*D*) between the Sender (S) and Receiver (R) states (*D_S_* – *D_R_*) as a function of λ_*J*_ for different values of λ_*N*_ at (d) λ_*D*_ = 10.0 and (e) λ_*D*_ = 20.0. All other parameters are standard.

**FIG. S5:**
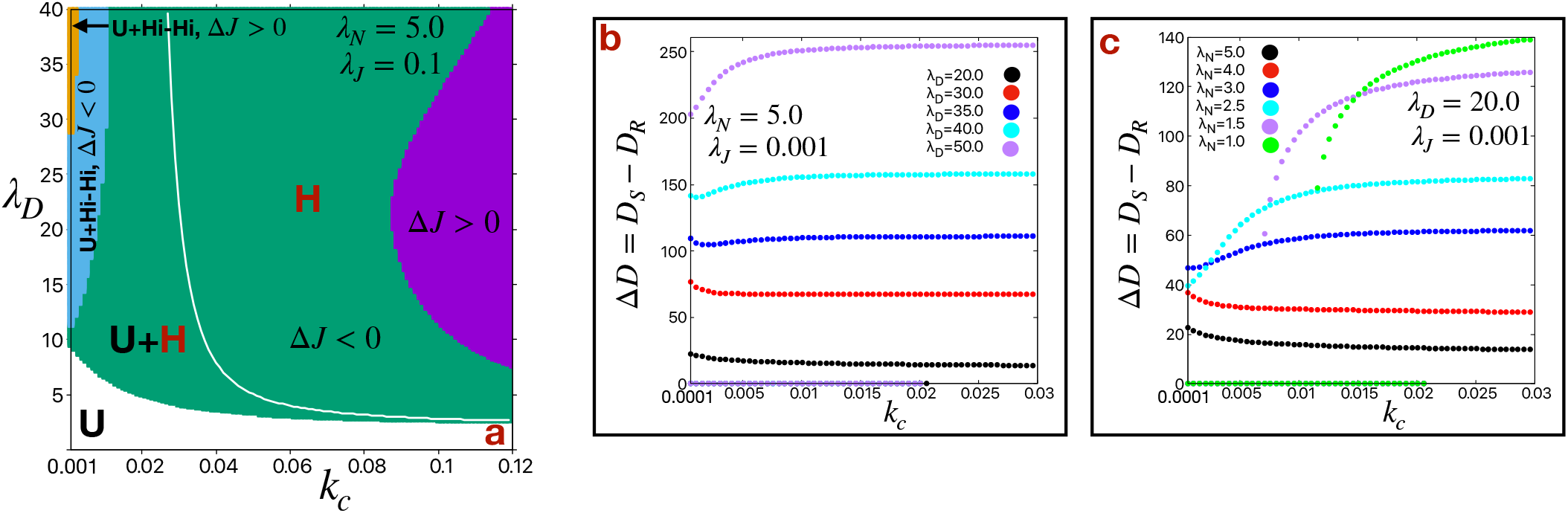
Effect of rate of cis-inhibition (*k_c_*). (a) Phase diagrams in λ_*D*_ – *k_c_* plane for λ_*N*_ = 5.0, λ_*J*_ = 0.1. The white and colored (green: Δ*J*(*J_S_* – *J_R_*) < 0) and purple: Δ*J*(*J_S_* – *J_R_*) > 0)) regions represent uniform (U: Δ*N*(*N_S_* – *N_R_*) = Δ*D*(*D_S_* – *D_R_*) = 0) and hexagon (H: Δ*N* < 0, Δ*D* > 0) phases respectively. The cyan (Δ*J* < 0) and orange (Δ*J* > 0) colored region represents the bistability of uniform (U) and High-High (Hi-Hi: Δ*N* > 0, Δ*D* > 0) phases. The white line represents the boundary of U region. The difference in Delta (Δ*D*) between the Sender (S) and Receiver (R) states (*D_S_* – *D_R_*) as a function of *k_c_* (b) at λ_*N*_ = 5.0, λ_*J*_ = 0.001 for different values of λ_*D*_ and (c) at λ_*D*_ = 20.0, λ_*J*_ = 0.001 for different values of λ_*N*_. All other parameters are standard.

**FIG. S6:**
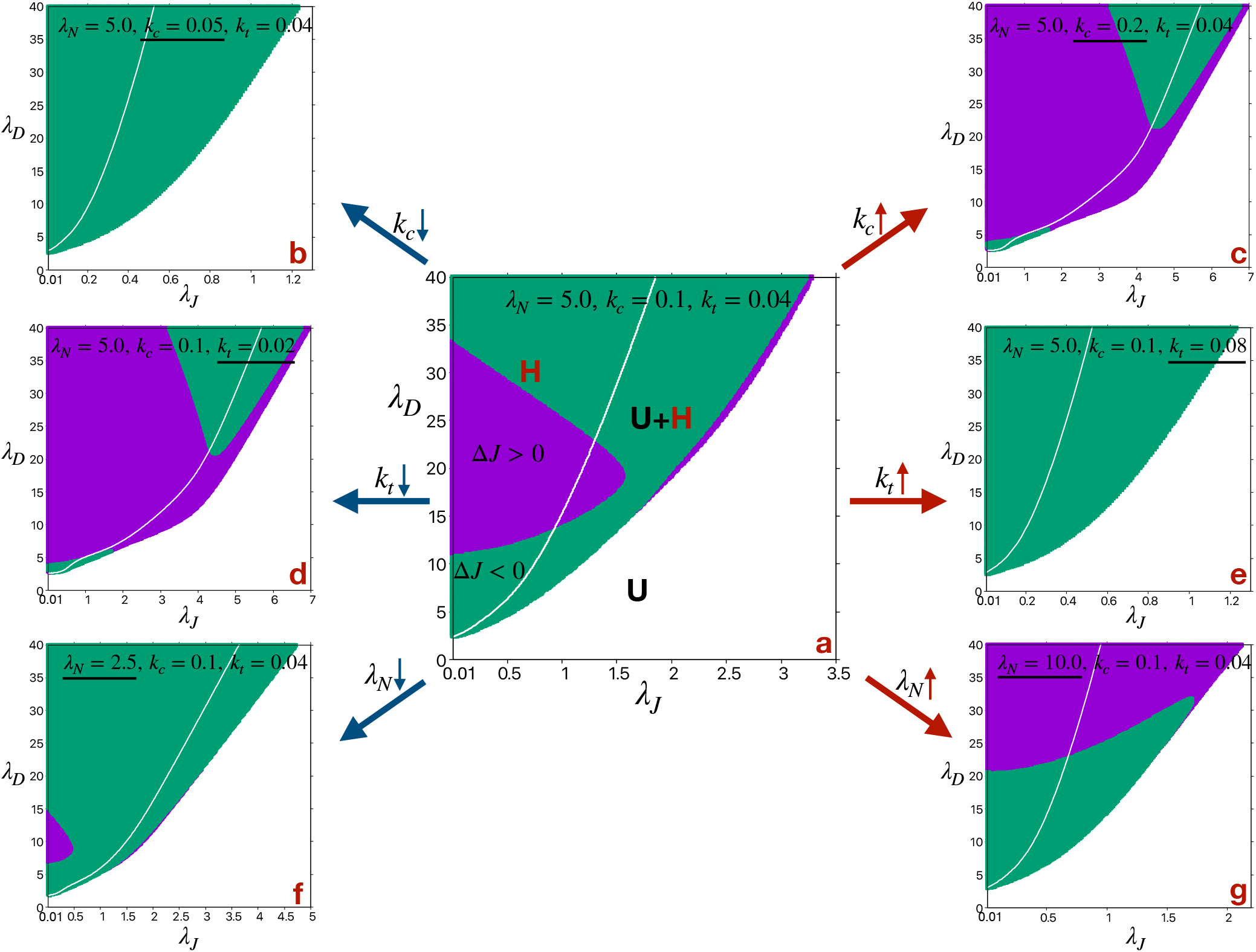
Change in phase diagram in λ_*D*_ – λ_*J*_ plane. The standard phase diagram (a) in λ_*D*_ – λ_*J*_ plane at λ_*N*_ = 5.0, *k_c_* = 0.1, *k_t_* = 0.04 changes, especially the region of bistability (where both the uniform (U: Δ*N*(*N_S_* – *N_R_*) = Δ*D*(*D_S_* – *D_R_*) = 0) and hexagon (H: Δ*N* < 0, Δ*D* > 0) phases are stable), with the decrease and increase in (b–c) *k_c_*, (d–e) *k_t_* and (f–g) λ_*N*_. The white and colored (green: Δ*J*(*J_S_* – *J_R_*) < 0) and purple: Δ*J*(*J_S_* – *J_R_*) > 0)) regions represent uniform (U) and hexagon (H) phases respectively. The white lines represent the boundary of U region. All other parameters are standard.

## Notes

### Competing Interest Statement

The authors have declared no competing interest.

