## Supplemental Information for "The Alternate Ligand Jagged Enhances the Robustness of Notch Signaling Patterns"

**The description of the video files**

Video S1: Spatiotemporal patterns of Delta ($D$) starting from uniform initial condition with small fluctuations for $\lambda_N = 5.0$, $\lambda_D = 10.0$, $\lambda_J = 0.5$ and $L = 6$. All other parameters are standard.

Video S2: Spatiotemporal patterns of Delta ($D$) starting from uniform initial condition with small fluctuations for $\lambda_N = 5.0$, $\lambda_D = 10.0$, $\lambda_J = 0.5$ and $L = 50$. All other parameters are standard.

Video S3: Spatiotemporal patterns of Delta ($D$) starting from uniform initial condition with small fluctuations for $\lambda_N = 1.5$, $\lambda_D = 10.0$ and $\lambda_J = 0.1$. All other parameters are standard.

Video S4: Spatiotemporal patterns of Delta ($D$) starting from a hexagonal seed for $\lambda_N = 5.0$, $\lambda_D = 10.0$ and $\lambda_J = 0.5$. All other parameters are standard.

Video S5: Spatiotemporal patterns of Delta ($D$) starting from a hexagonal seed for $\lambda_N = 5.0$, $\lambda_D = 10.0$ and $\lambda_J = 0.9$. All other parameters are standard.

**The reduced set of 12 ODE's**

$$\dot{N}_A = \lambda_N(1 + I_A^{n_N}/(1 + I_A^{n_N})) - N_A(k_c(D_A + J_A) + 0.5k_t(D_B + D_C + J_B + J_C)) - \gamma N_A$$
$$\dot{N}_B = \lambda_N(1 + I_B^{n_N}/(1 + I_B^{n_N})) - N_B(k_c(D_B + J_B) + 0.5k_t(D_C + D_A + J_C + J_A)) - \gamma N_B$$
$$\dot{N}_C = \lambda_N(1 + I_C^{n_N}/(1 + I_C^{n_N})) - N_C(k_c(D_C + J_C) + 0.5k_t(D_A + D_B + J_A + J_B)) - \gamma N_C$$
$$\dot{D}_A = \lambda_D/(1 + I_A^{n_D}) - D_A(k_c N_A + 0.5k_t(N_B + N_C)) - \gamma D_A$$
$$\dot{D}_B = \lambda_D/(1 + I_B^{n_D}) - D_B(k_c N_B + 0.5k_t(N_C + N_A)) - \gamma D_B$$
$$\dot{D}_C = \lambda_D/(1 + I_C^{n_D}) - D_C(k_c N_C + 0.5k_t(N_A + N_B)) - \gamma D_C$$
$$\dot{J}_A = \lambda_J(1 + I_A^{n_J}/(1 + I_A^{n_J})) - J_A(k_c N_A + 0.5k_t(N_B + N_C)) - \gamma J_A$$
$$\dot{J}_B = \lambda_J(1 + I_B^{n_J}/(1 + I_B^{n_J})) - J_B(k_c N_B + 0.5k_t(N_C + N_A)) - \gamma J_B$$
$$\dot{J}_C = \lambda_J(1 + I_C^{n_J}/(1 + I_C^{n_J})) - J_C(k_c N_C + 0.5k_t(N_A + N_B)) - \gamma J_C$$
$$\dot{I}_A = 0.5k_t N_A(D_B + D_C + J_B + J_C) - \gamma I_A$$
$$\dot{I}_B = 0.5k_t N_B(D_C + D_A + J_C + J_A) - \gamma I_B$$
$$\dot{I}_C = 0.5k_t N_C(D_A + D_B + J_A + J_B) - \gamma I_C$$

| Parameters | | | | | |
|---|---|---|---|---|---|
| Figures | $\lambda_N$ | $\lambda_D$ | $\lambda_J$ | $k_c$ | $k_t$ |
| Figure 2 | 1.822, 0.9, 5.0 | [1.0 – 4.0] | 0.1 | 0.1 | 0.04 |
| Figure 3 | [0.5 – 3.0] | [0.01 – 40.0] | [0.01 – 3.5] | 0.1 | 0.04 |
| Figure 4 | 5.0 | 10.0, 20.0 | 0.001, 0.5, 0.65 | 0.1 | 0.04 |
| Figure 5 | 5.0 | 5.0, 10.0, 20.0, 40.0 | [0.001 – 10] | 0.1 | 0.04 |
| Figure 6 | 5.0 | 20.0 | [0.001 – 4.5] | [0.001 – 0.12] | [0.01 – 0.09] |
| Figure 7 | 5.0 | 10.0 | 0.5, 0.9 | 0.1 | 0.04 |
| Figure S1 | 5.0 | 10.0 | 0.001, 0.5 | 0.1 | 0.04 |
| Figure S2 | 5.0 | 5.0, 10.0, 20.0 | 0.001, 0.5 | 0.1 | 0.04 |
| Figure S3 | [1.5 – 3.0] | 10.0 | [0.1 – 1.3] | 0.1 | 0.04 |
| Figure S4 | [1.5 – 5.0] | [5.0 – 40.0] | [0.001 – 10] | 0.1 | 0.04 |
| Figure S5 | [1.0 – 5.0] | [0.01 – 50.0] | 0.001, 0.1 | [0.001 – 0.12] | 0.04 |
| Figure S6 | 2.5, 5.0, 10.0 | [0.01 – 40.0] | [0.01 – 7] | 0.05, 0.1, 0.2 | 0.02, 0.04, 0.08 |

TABLE S1: **Parameters for all the Figures described in the manuscript.**

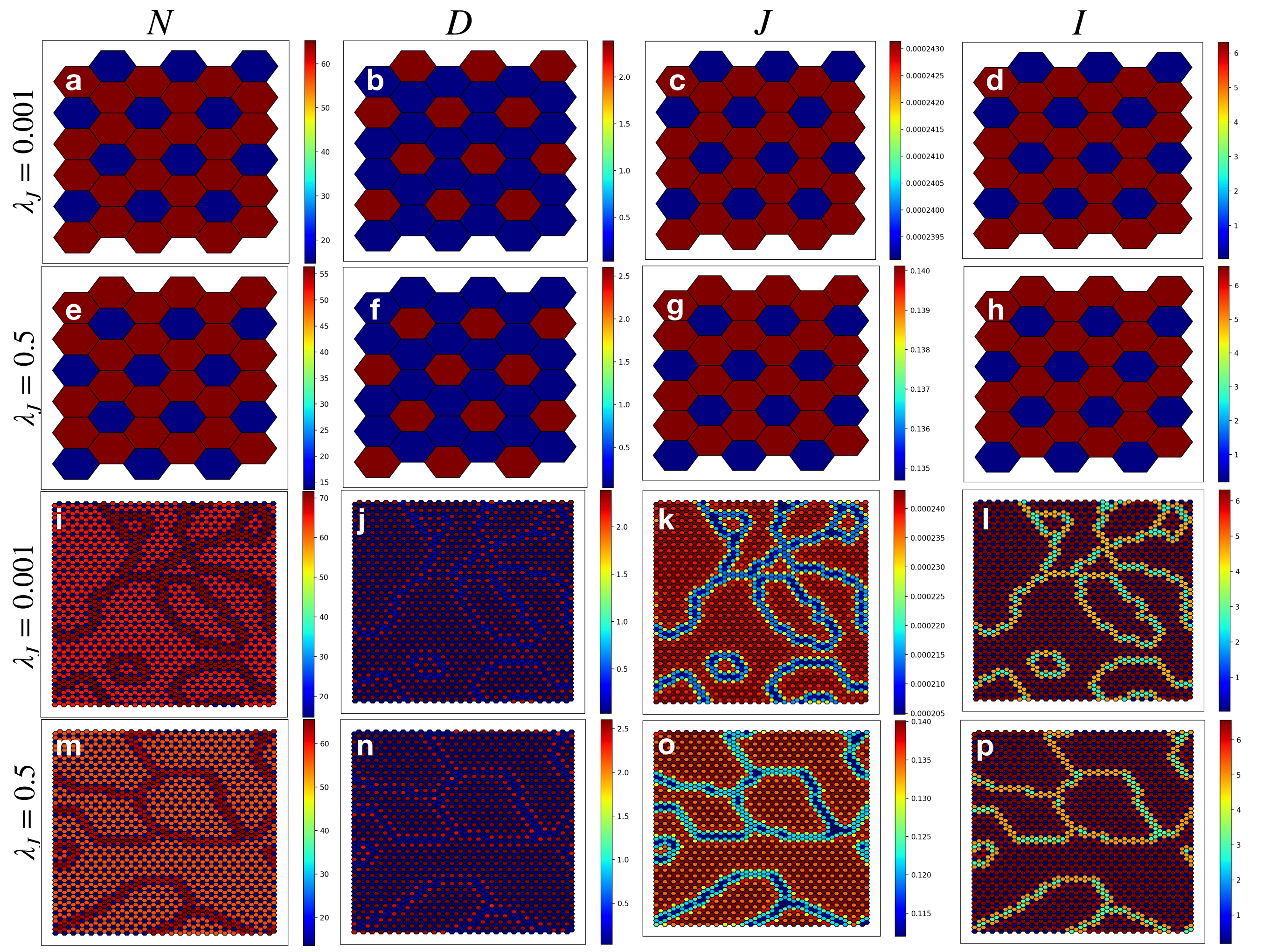


FIG. S1: **Steady state patterns of Notch (N), Delta (D), Jagged (J) and NICD (I) on a hexagonal lattice.** The steady states of Notch (N), Delta (D), Jagged (J) and NICD (I) at $\lambda_N = 5.0$, $\lambda_D = 10.0$ and $\lambda_J = 0.001$ and 0.5 on a hexagonal lattice of size (a–h) $L = 6$ and (i–p) $L = 50$. All other parameters are standard.

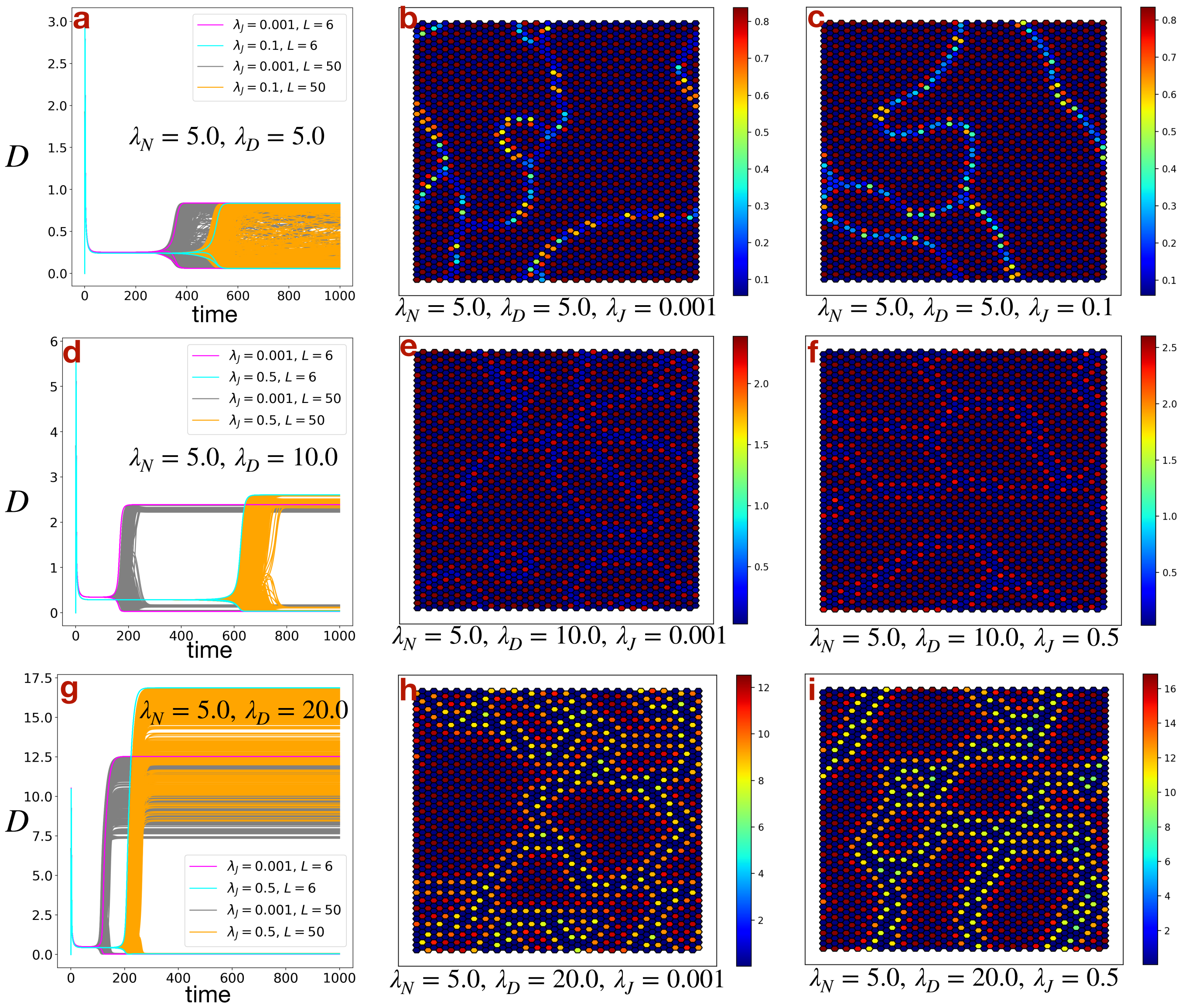


FIG. S2: **Effect of production rate of Delta ($\lambda_D$).** Dynamics of Delta (D) for all the cells on a hexagonal lattice of size $L = 6$ and $L = 50$ at $\lambda_N = 5.0$, $\lambda_J = 0.001$ and 0.5 (0.1 instead of 0.5 for $\lambda_D = 5.0$, because at $\lambda_J = 0.5$, $\lambda_D = 5.0$ and $\lambda_N = 5.0$, the steady states become uniform (U)) for (a) $\lambda_D = 5.0$, (d) $\lambda_D = 10.0$ and (g) $\lambda_D = 20.0$. (b), (c), (e), (f), (h), (i) Corresponding patterns of Delta (D) at steady states for $L = 50$ for different values of $\lambda_D$ and $\lambda_J$. All other parameters are standard.

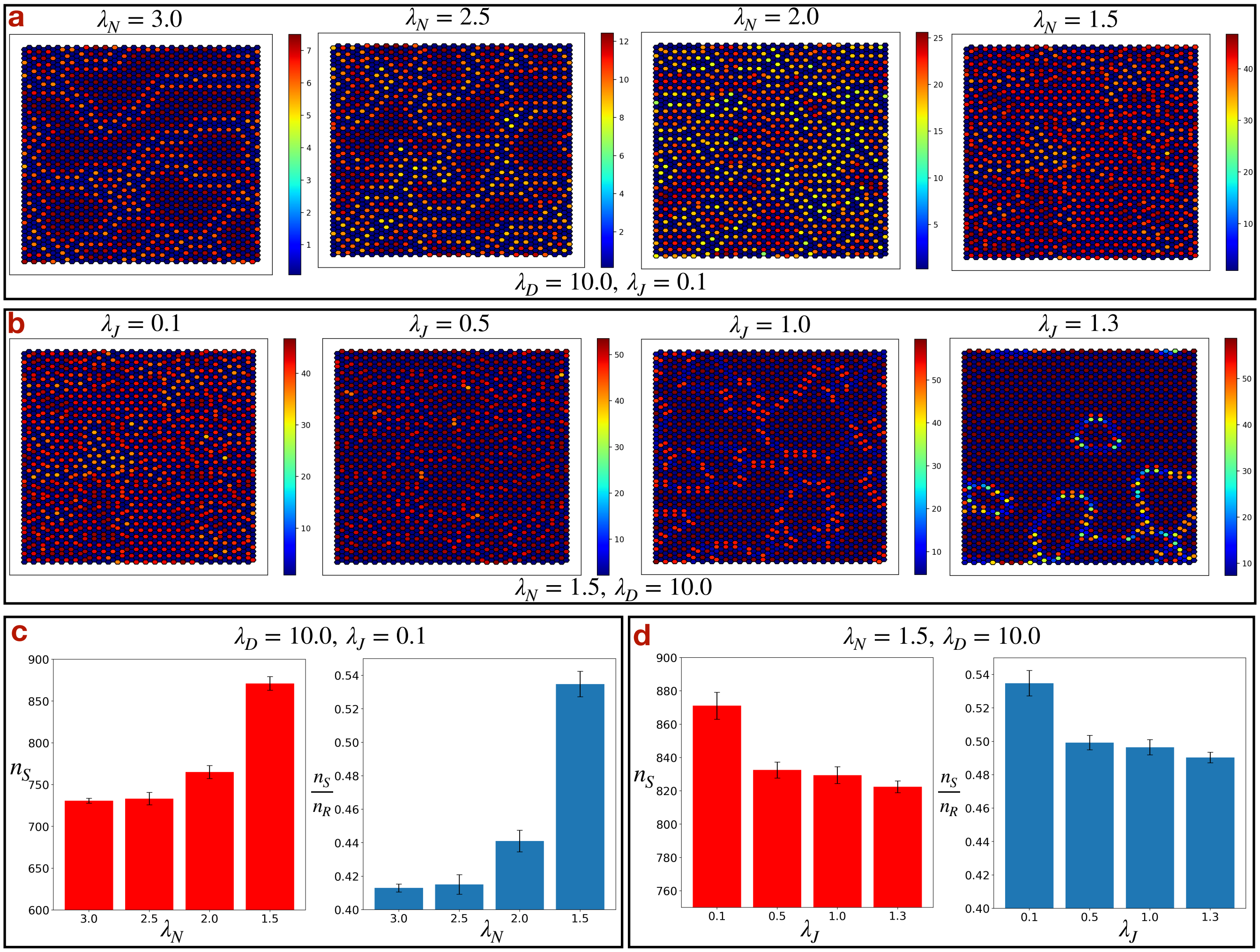


FIG. S3: **Effect of production rate of Notch ($\lambda_N$).** The steady state patterns of Delta ($D$) on a hexagonal lattice of size $L = 50$ (a) for varying $\lambda_N$ at $\lambda_D = 10.0$, $\lambda_J = 0.1$ and (b) for varying $\lambda_J$ at $\lambda_N = 1.5$, $\lambda_D = 10.0$. (c), (d) The corresponding number of Sender (S) states ($n_S$) and ratio of number of Sender (S) and Receiver (R) states ($\frac{n_S}{n_R}$). All other parameters are standard.

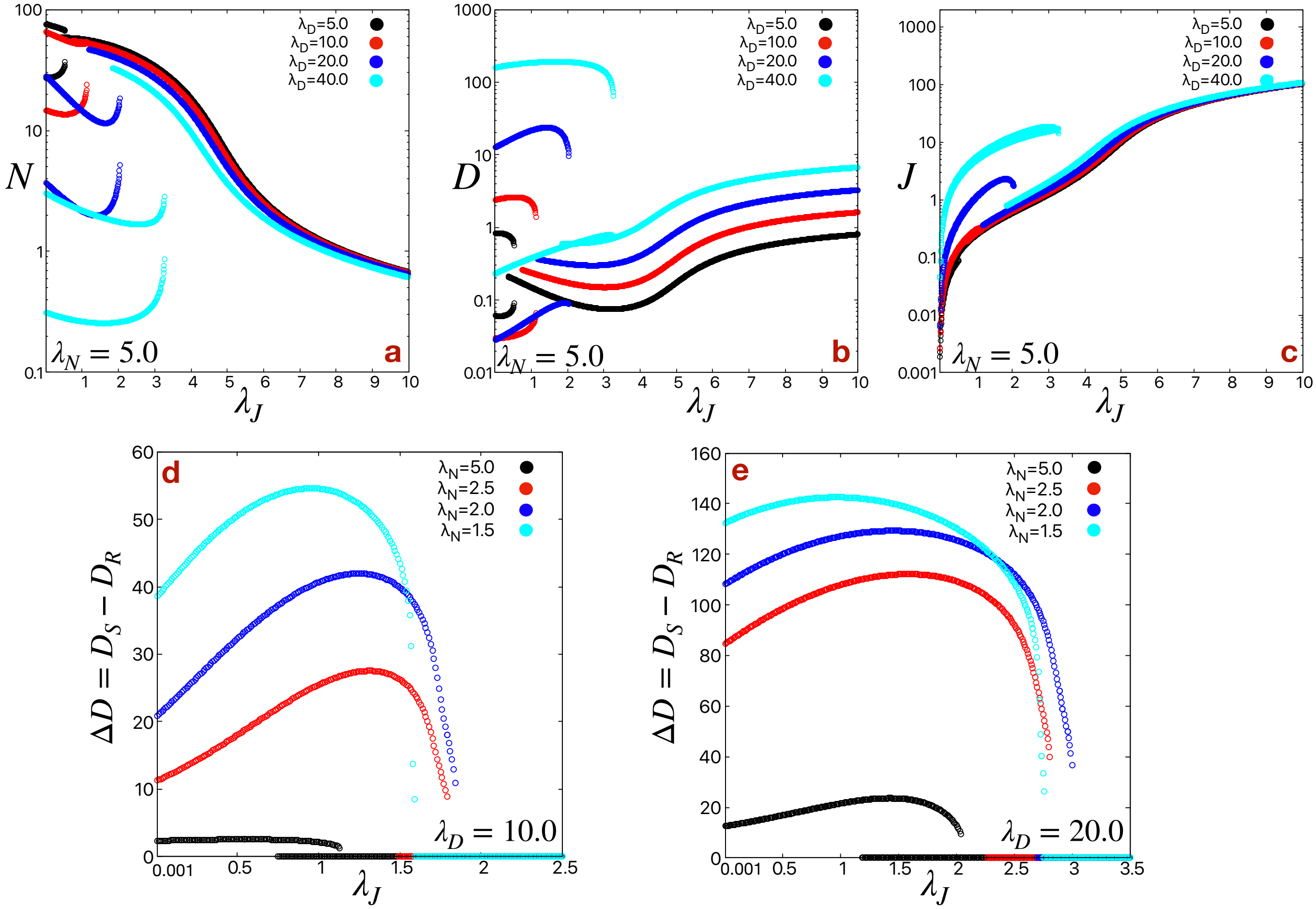


FIG. S4: **Steady state values of Notch (N), Delta (D) and Jagged (J) as a function of $\lambda_J$.** Steady state values of (a) Notch (N), (b) Delta (D), (c) Jagged (J) as a function of $\lambda_J$ for different values of $\lambda_D$ at $\lambda_N = 5.0$. The y-axes ($N$, $D$ and $J$-axes) are in log-scale. The difference in Delta ($\Delta D$) between the Sender (S) and Receiver (R) states ($D_S - D_R$) as a function of $\lambda_J$ for different values of $\lambda_N$ at (d) $\lambda_D = 10.0$ and (e) $\lambda_D = 20.0$. All other parameters are standard.

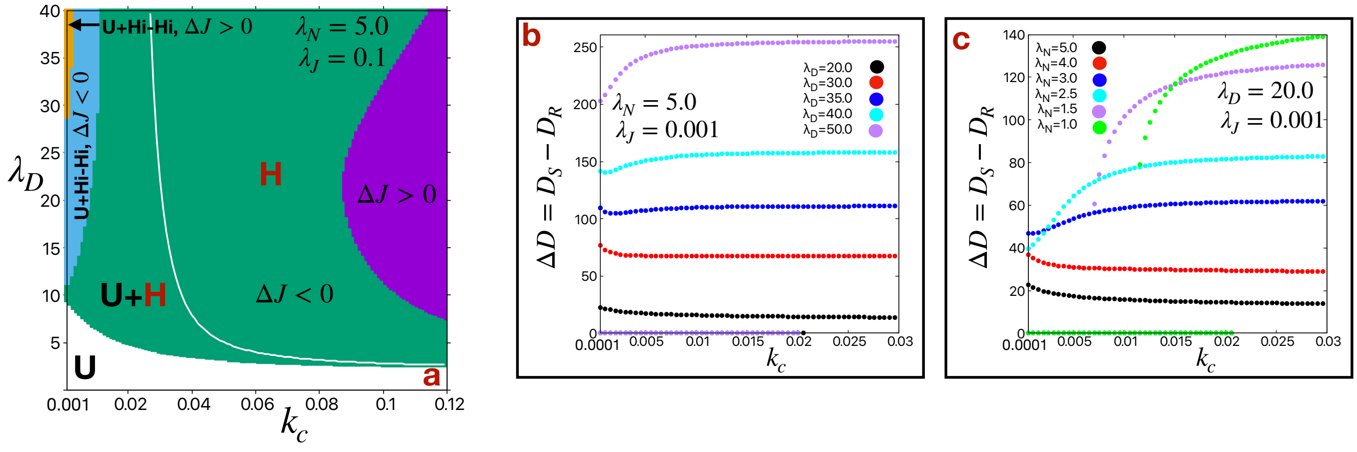


FIG. S5: **Effect of rate of cis-inhibition ($k_c$).** (a) Phase diagrams in $\lambda_D - k_c$ plane for $\lambda_N = 5.0$, $\lambda_J = 0.1$. The white and colored (green: $\Delta J(J_S - J_R) < 0$) and purple: $\Delta J(J_S - J_R) > 0$)) regions represent uniform (U: $\Delta N(N_S - N_R) = \Delta D(D_S - D_R) = 0$) and hexagon (H: $\Delta N < 0$, $\Delta D > 0$) phases respectively. The cyan ($\Delta J < 0$) and orange ($\Delta J > 0$) colored region represents the bistability of uniform (U) and High-High (Hi-Hi: $\Delta N > 0$, $\Delta D > 0$) phases. The white line represents the boundary of U region. The difference in Delta ($\Delta D$) between the Sender (S) and Receiver (R) states ($D_S - D_R$) as a function of $k_c$ (b) at $\lambda_N = 5.0$, $\lambda_J = 0.001$ for different values of $\lambda_D$ and (c) at $\lambda_D = 20.0$, $\lambda_J = 0.001$ for different values of $\lambda_N$. All other parameters are standard.

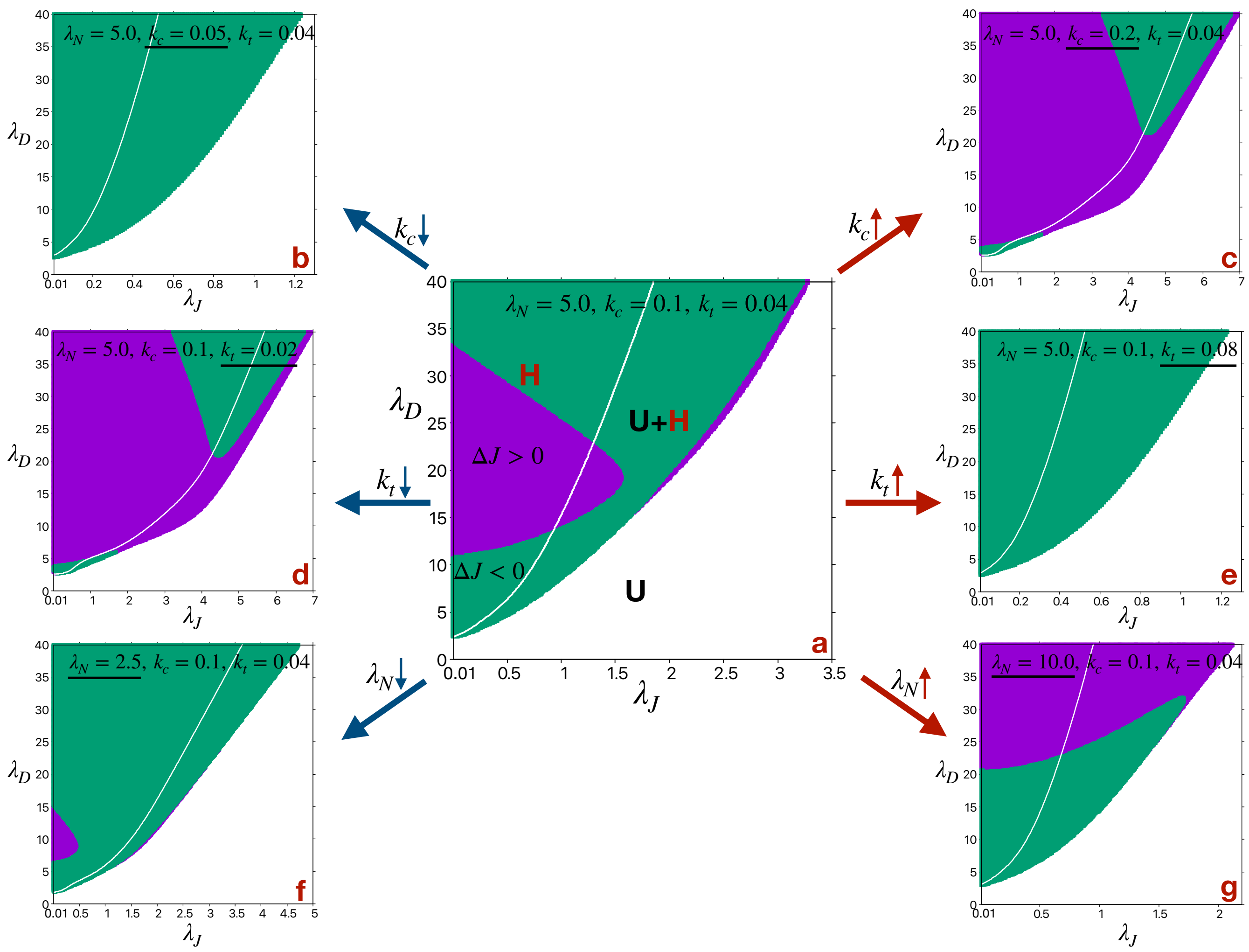


FIG. S6: **Change in phase diagram in $\lambda_D - \lambda_J$ plane.** The standard phase diagram (a) in $\lambda_D - \lambda_J$ plane at $\lambda_N = 5.0$, $k_c = 0.1$, $k_t = 0.04$ changes, especially the region of bistability (where both the uniform (U: $\Delta N(N_S - N_R) = \Delta D(D_S - D_R) = 0$) and hexagon (H: $\Delta N < 0$, $\Delta D > 0$) phases are stable), with the decrease and increase in (b–c) $k_c$, (d–e) $k_t$ and (f–g) $\lambda_N$. The white and colored (green: $\Delta J(J_S - J_R) < 0$) and purple: $\Delta J(J_S - J_R) > 0$)) regions represent uniform (U) and hexagon (H) phases respectively. The white lines represent the boundary of U region. All other parameters are standard.
